# Actin polymerization counteracts prewetting of N-WASP on supported lipid bilayers

**DOI:** 10.1101/2024.04.14.589463

**Authors:** Tina Wiegand, Jinghui Liu, Anatol W. Fritsch, Lutz Vogeley, Isabel LuValle-Burke, Jan Geisler, Anthony A. Hyman, Stephan W. Grill

**Author notes:** These authors contributed equally (Tina Wiegand, Jinghui Liu).

## Abstract

Cortical condensates, transient punctate-like structures rich in actin and the actin nucleation pathway member N-WASP, form during activation of the actin cortex in the *C. elegans* oocyte. Their emergence and spontaneous dissolution is linked to a phase separation process driven by chemical kinetics. However, the physical process that drives the onset of cortical condensate formation near membranes remains unexplored. Here, using a reconstituted phase separation assay of cortical condensate proteins, we demonstrate that the key component, N-WASP, can collectively undergo surface condensation on supported lipid bilayers via a prewetting transition. Actin partitions into the condensates, where it polymerizes and counteracts the N-WASP prewetting transition. Taken together, the dynamics of condensate-assisted cortex formation appear to be controlled by a balance between surface-assisted condensate formation and polymer-driven condensate dissolution. This opens new perspectives for understanding how the formation of complex intracellular structures is affected and controlled by phase separation.

## Introduction

Surface transitions of biomolecules near membranes can serve as an important mechanism to localize biochemical reactions in the cell. The affinity of proteins to membranes can induce their phase separation on the membrane (1–4), far below the saturation concentration for bulk liquid-liquid phase separation (LLPS) (5, 6). This process, termed prewetting, is related to wetting phenomena (7–9). How prewetting condensates form on membranes remains poorly understood. Furthermore, understanding wetting phenomena in biological systems that typically are far from equilibrium remains challenging. While the principles of LLPS are still applicable within localized regions within cells that remain equilibrated (10), chemical reactions can drive condensates out of equilibrium and significantly influence phase separation kinetics (11–13). Condensates can be dynamic in nature, and recruit additional components (14) that drive specific biochemical reactions, such as actin polymerization (2, 15–17), which can even lead to their dissolution (18). Examples for dynamic condensates forming in vivo are the actin fusion focus in budding yeast (19), and cortical condensates that transiently form prior to their self-organized dissassembly at the oocyte to em-bryo transition (20).

These two examples indicate that condensates that form in the cytoskeletal context can be highly dynamic. The activation of the actomyosin cortex in the *C. elegans* embryo is accompanied by the transient formation of hundreds of punctate-like objects, termed cortical condensates, that are rich in the actin nucleation pathway member N-WASP, the branched nucleator Arp2/3, and actin (20). Cortical condensates exhibit chemical dynamics that obey mass-action kinetics to control composition and size, resulting in a dynamic instability that orchestrates a switch from condensate growth to shrinkage and dissolution. The molecular mechanisms that are at the heart of this switch remain unclear. Furthermore, N-WASP has been shown to undergo LLPS in vitro together with its adapter proteins NCK and Nephrin, but it remains unclear how this behaviour impacts on cortical condensate formation (21, 22).

Here we set out to understand the mechanisms that drive condensation of cortical condensates and that orchestrate their switch to disassembly. We employ an in vitro approach to investigate the formation of liquid-like condensates on supported lipid bilayers (SLBs) in order to reveal the spatiotemporal dynamics of key proteins that constitute cortical condensates dynamics. Our study encompasses two variants of N-WASP, human and *C. elegans*, in order to allow for comparisons with previously published findings, both in human and *C. elegans* systems.

## Results

### N-WASP condenses in absence of adaptors in vitro

In the *C. elegans* oocyte the adaptor proteins Nck and ephrin are dispensable for the formation of cortical condensates (20). We first asked if homotypic-interactions between N-WASP proteins are sufficient to drive N-WASP condensation. We purified full length N-WASP, both from human and *C. elegans*, recombinantly and performed phase separation assays in vitro (for quality controls of different protein constructs and activity assay towards actin polymerization see Fig. 1a and Fig. S1a-c). We find that purified full length *C. elegans* N-WASP undergoes phase separation at physiological salt concentrations (150 mM KCl) in vitro without further adaptor proteins or crowding agents (Fig. 1d and Fig. S1d). The critical concentration for phase separation in bulk solution is *∼*500 nM, as determined by the volume fraction of the condensed phase in bulk (Fig. S1e). However, typical concentrations of N-WASP in *C. elegans* embryos are *∼*100 nM (23), indicating that bulk phase separation of N-WASP should not take place in vivo. We next focused on human N-WASP (Fig. S2a-b), which similarly undergoes LLPS without additional proteins. We determined the associated saturation concentration to be *∼*1 *μ*M (Fig. S2c). We conclude that homotypic interactions are sufficient to drive bulk condensation of N-WASP.

**Fig. 1.**
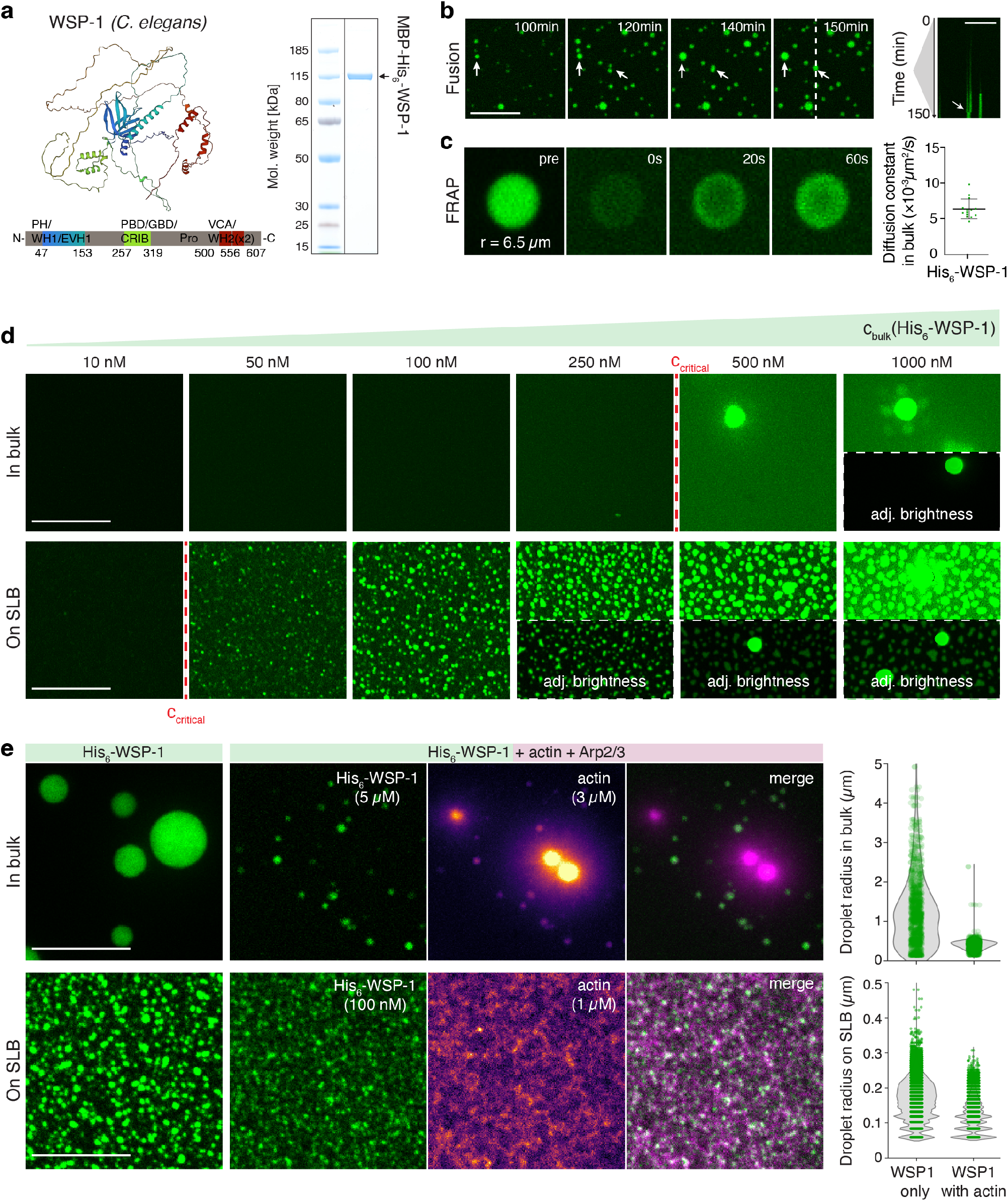
Actin controls equilibrium size of N-WASP condensates in vitro. a) Alpha fold predicts large disordered regions for *C. elegans* N-WASP (WSP-1) (sequence dependent color code). Sodium dodecyl sulphate gel shows recombinantly expressed and purified MBP-His6-WSP-1 (for activity controls and results of human N-WASP see SI). b) Confocal time-lapse of His6-WSP-1 (5 *μ*M, 10 % 488-tagged, MBP-tag cleaved right before experiment, see SI) forming droplets in bulk upon lowering KCl concentration to 150 mM. Kymograph indicates coarsening of droplets and fusion (white arrows). c) Fluorescence recovery after photo-bleaching (FRAP) images of condensate formed from 5 *μ*M His_6_-WSP-1 between 5 to 60 min. Diffusion coefficients D inside the condensates were determined for n = 14 condensates. Data are the mean ± s.d. d) Phase separation assays of His_6_-WSP-1 reveals a critical concentration for phase separation of 500 nM in bulk (upper row, see SI Figure S1e) and 50 nM on a supported lipid bilayer (SLB) with 1 % Ni-NTA lipids (lower row). e) Left: Maximum projections of confocal z-stacks of His_6_-WSP-1 condensates formed over 1 h in bulk (upper row, see Movie S3) and on a SLB (lower row, see Movie S4) in actin polymerization buffer alone (left images), and in the presence of actin (10 % AF647 labeled, magenta) and Arp2/3. Right: Quantification of droplet radii from max projections. All scale bars 10 *μ*m.

### N-WASP in vitro condensates show liquid features

In vivo cortical condensates were found to form via a mechanism different from equilibrium LLPS, but display characteristics indicative of rapid internal mixing (20). Do in vitro N-WASP condensates also show liquid like features? We performed time-lapse imaging and observed coarsening and fusion (Fig. 1b and Movie S1) of in vitro condensates of *C. elegans* N-WASP. We further conducted FRAP experiments showing exchange of molecules between the condensates and the surrounding dilute phase (Fig. 1c) and determined the diffusion rate within the condensates to be 0.006 *±* 0.001 *μ*m^2^/s (mean *±* s.d., Fig. 1d). Similarly, human N-WASP in vitro condensates also exchange material with the bulk. Compared to *C. elegans* N-WASP, we observed the diffusion coefficient of N-WASP inside the condensates to be higher (0.098 *±* 0.036 *μ*m^2^/s; Fig. S2d). We conclude that N-WASP condensates show liquid features such as coalescence, fusion and diffusion, albeit with slow dynamics.

### N-WASP forms condensates on SLBs at physiological concentration

Typical in vivo concentrations of N-WASP are below the critical concentration (c_sat_) of bulk condensation of N-WASP (Fig. S1e). Do interactions with membranes allow for N-WASP condensation below c_sat_? We purified His-tagged N-WASP and performed phase separation assays in the presence of SLBs. The surface interaction strength was adjusted by tuning the concentration of DGS-Ni-NTA within the SLB. At physiological N-WASP concentration (100 nM), dense objects appear to form on SLBs at 0.5 % DGS-Ni-NTA (Fig. S3a). We used 1 % DGS-Ni-NTA for subsequent experiments, as the condensates resemble in vivo appearance and show liquid features (Fig. S3b and Movie S2). At 1 % DGS-Ni-NTA content condensates start forming on the membrane at bulk His-N-WASP concentrations of 50 nM (Fig. 1b lower row, Fig. S3d). We asked if these objects are monolayered or multiplayered structures. If they were monolayered, their integrated fluorescence intensity should scale linearly with covered area on the SLB. If they were multilayered, the integrated N-WASP fluorescence intensity should scale with covered area with an exponent greater than 1 (see SI). We find that at 100 nM bulk concentration, this scaling exponent is above 1: 1.10 *±* 0.06 for *C. elegans* and 1.12 *±* 0.04 for human N-WASP (Fig. S3c and S4, also see SI section on multilayered condensates). We conclude that N-WASP can form multilayered surface condensates at concentrations significant lower than the critical concentration c_sat_ for bulk phase separation.

### Actin limits final size of in vitro N-WASP condensates

In the *C. elegans* embryo, actin accumulates in cortical N-WASP condensate over time. This results in negative feedback onto N-WASP and a dynamic instability via a self-organized switch to cortical condensate disassembly (20). Do in vitro N-WASP condensates nucleate the growth of actin fibers, and do these then limit N-WASP condensate growth? We set out to answer this question separately in bulk and on SLBs. First, we added actin and Arp2/3 during the formation of bulk *C. elegans* N-WASP condensates under polymerizing conditions (buffer contained 0.1 mM Mg^2+^ and 0.2 mM ATP). We find that actin monomers concentrate in N-WASP condensates, and that actin fibres polymerize and grow out of the condensates (Fig. 1e and SI Movie 3). At later times some condensates appear to disassemble, leaving behind dense actin networks. After 1 hr, we find condensates formed in the presence of actin to be of a smaller size on average (0.4 *±* 0.1 *μ*m in radius) compared to those formed without actin (1.2 *±* 0.9 *μ*m in radius) (Fig. 1e upper row). We conclude that actin filaments limit the growth of bulk N-WASP condensates. Next, we investigated if surface condensation of *C. elegans* N-WASP on SLBs is impacted in a similar way by actin. We find that on SLBs condensates are reduced in size in the presence of actin & Arp2/3 (0.12 *±* 0.04 *μ*m compared to 0.16 *±* 0.06 *μ*m without actin in radius) (Fig. 1e lower row and SI Movie 4). We observed similar effects of actin on human N-WASP surface condensates (Fig. S5). We conclude that N-WASP condensates concentrate actin monomers and serve as nucleation hubs for actin polymerization. In addition, actin fibres appear to disassemble N-WASP condensates and thereby limit their final size, similar to what is seen in the cortical condensates in the *C. elegans* zygote.

### Actin polymerization feeds back on the growth of N-WASP condensates in an ATP-independent manner

We next set out to understand if the negative feedback mechanism of actin on condensate growth is initiated by monomeric actin, or by the polymerized actin meshwork inside the cortical condensates. Is this negative feedback dependent on ATP consumption? To answer these questions, we added actin monomers without Arp2/3 or in the presence of polymerization inhibiting drugs to the N-WASP condensates. Actin still polymerizes into actin filaments and limits the size of N-WASP condensates even in the absence of Arp2/3, however, condensates are bigger (r = 0.6 *±* 0.2 *μ*m) than in the presence of Arp2/3 (r = 0.4 *±* 0.1 *μ*m)(Fig. S6a). Suppressing actin polymerization by latrunculin A gives rise to bulk condensates that are still enriched with actin, but that have a size (1.4 *±* 0.9 *μ*m) that is not significantly different from condensates formed in the absence of latrunculin A (Fig. S6a). Together, this suggests that polymerized actin is responsible for limiting the size of N-WASP condensates. While actin polymerization is an active process, not all steps require ATP (24, 25). Filament elongation is ATP-independent, and ATP molecules subsequently undergo hydrolysis within the actin filament (26). Likewise, ATP bound to Arp2/3 is hydrolysed following filament branching (27). We next evaluated if the size-limiting effect on N-WASP condensates by actin polymerization depends on ATP hydrolysis. To this end we analysed the growth of actin bound with the non-hydrolyzing ATP variant AMP-PNP in ATP free buffer and determined the effect on N-WASP condensation. We find that the kinetics of actin polymerization of nucleotide-exchanged actin are slowed down, but do not completely stall (Fig. S6b). The final size of N-WASP condensates in the presence of nucleotide-exchanged actin (r = 0.4 *±* 0.2 *μ*m) is similar compared to the ATP rich conditions (Fig. S6c). We conclude that N-WASP condensate size is limited by the N-WASP dependent polymerizaton of actin filaments in a manner that is independent of ATP.

### Wetting phenomena of condensates on biological surfaces

We note that the presence of a surface plays an important role for the condensation of N-WASP at physiological concentrations. Next, we sought to understand the mechanism by which N-WASP forms multilayered surface-associated condensates. Interactions between condensates and surfaces fall into the category of wetting phenomena (7). Above the critical saturation concentration for liquid-liquid phase separation, condensates formed in bulk can wet biological surfaces (Fig. 2a-b, III). As a result of affinity-induced adsorption, proteins can have increased concentrations near biological surfaces (Fig. 2a-b, I). Therefore, surface condensates can also form below the saturation concentration via a prewetting transition (Fig. 2a-b, II). As a type of critical phenomena, prewetting transitions were reported for sequence-dependent protein condensation processes on DNA (28). For lipid membranes, it has been postulated that prewetting transitions could also play a role for protein surface condensation (9, 29, 30).

**Fig. 2.**
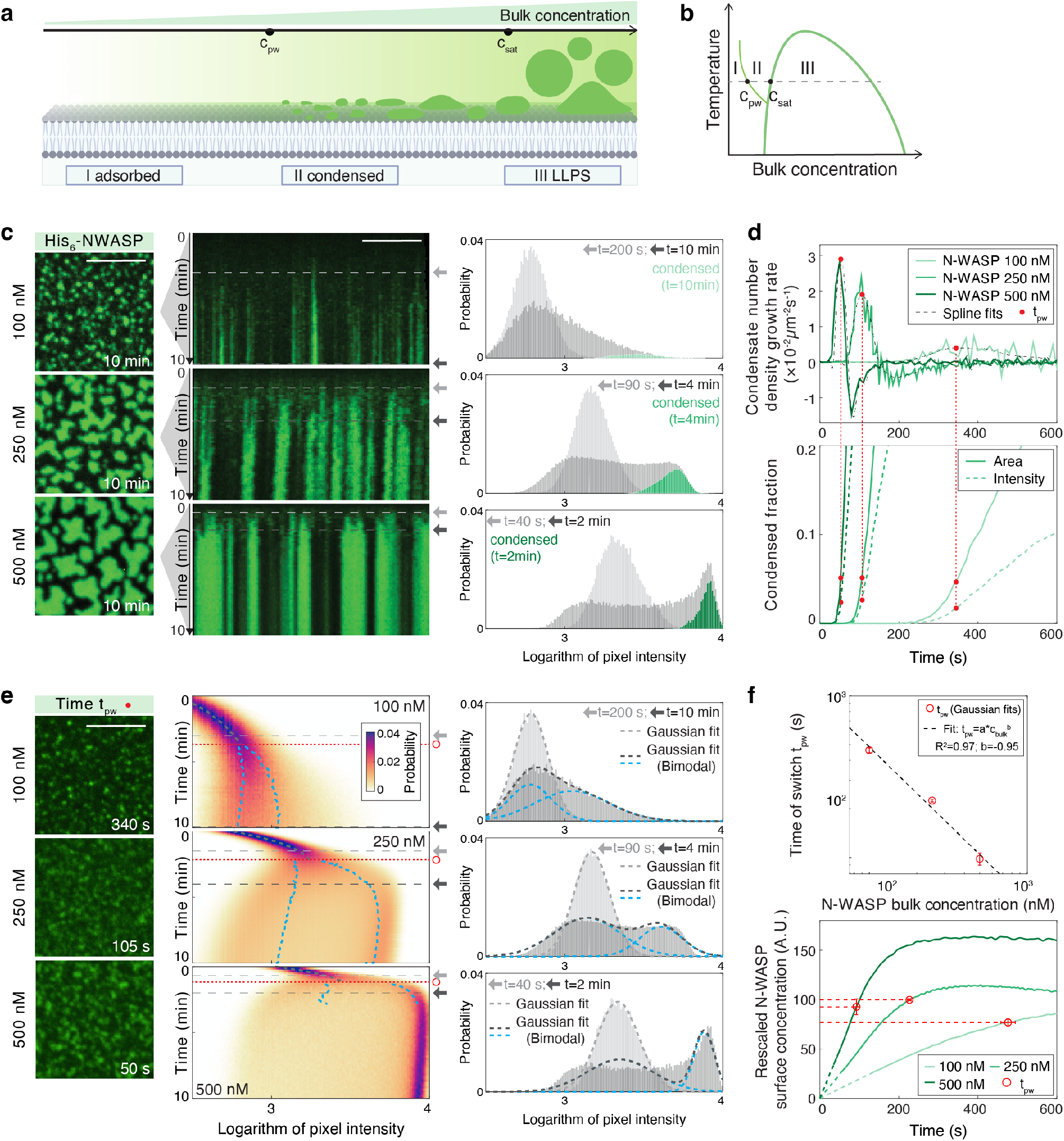
N-WASP prewets on supported lipid bilayers. a) Proteins interactions with membranes depends on their bulk concentration. At low concentrations, proteins adsorb as a thin layer (I, left). Crossing the critical concentration for prewetting (c_*p*_*w*) condensed areas form (II, middle). Above the saturation concentration (c_*s*_*at*) for liquid-liquid phase separation (III, LLPS) droplets form spontaneously in bulk (right). b) Phase diagram depicting these three regions of a binary fluid in the presence of a surface. Prewetting line discriminates between areas of adsorption and surface condensation (prewetting). c) Confocal images of human His6-N-WASP (10 % AF488 labeled, 100 nM upper row, 250 nM middle, 500 M low) condensates on SLBs after t = 10 min. Scale bar = 10 *μ*m. Kymographs show adsorption and condensation phase over 10 min (SI Movie S5). Histogram of the logarithm of pixel intensities (right) shows enrichment of the condensed phase over time. Two distinct time points marked with light gray and dark gray arrows are chosen respectively for each time series. d) Condensate number density growth rate (upper) and condensed phase area (lower, solid lines) and intensity (lower, dashed lines) fractions quantified as a function of time from c). Red filled circles show the time when maximum condensation rate is reached. e) Left: Confocal images at the time point (red filled circles) identified in d). Middle: Probability density kymographs of the logarithm of pixel intensities for time-lapse images from c). Gray lines (single Gaussian) and blue lines (sum-of-two-Gaussians) represent overlays of probability density fits from either method. The time point of high- and low-intensity-branch splitting (red open circles) is determined by comparing fitting residues between the two methods. Right: Histograms and fits (grey dashed lines) for the 3 different N-WASP bulk concentrations each at 2 distinct time points (arrows). In the case of a sum-of-two-Gaussians fit, blue dashed lines show respectively the fitted lower and higher Gaussian peaks. f) Upper: The time of abrupt switch inversely scales with initial bulk concentrations. Error bars represent the range extracted from 10% variation of the threshold used for determining *t*_*pw*_ . For segmented images, the threshold is the growth rate of condensate number density; For splits of probability density branches, the threshold is the residue difference between single- and sum-of-two Gaussian fits (SI). Lower: The curated (SI) mean pixel intensity at the time of abrupt switch is comparable across different initial bulk concentrations.

### Abrupt N-WASP surface condensation following N-WASP adsorption

In our in vitro experiments, do N-WASP proteins undergo a prewetting transition on the supported lipid bilayer surface? The prewetting transition is discontinuous, and the height of the surface layer is predicted to increase abruptly as bulk concentration increases beyond the prewetting concentration (31). We anticipate to observe signatures of this critical behaviour at the onset of in vitro cortical condensate formation, as seen for example for prewetting condensate formation of transcription factors on DNA (28). There, histograms of surface protein amounts underwent a split up into an absorbed layer with low protein amounts and a condensed layer with higher protein amounts with time post adsorption of proteins to the DNA surface. To check for signatures of a prewetting transition for our in vitro cortical condensates, we set out to examine the kinetic process of N-WASP accumulation on supported lipid bilayers (Fig. 2c, Fig. S7a-b). We focused on the human N-WASP protein because of its higher mobility in bulk condensates and its reduced propensity of hardening (Fig. S2d, compared to the *C. elegans* variant Fig. 1c; data not shown). At time *t* = 0, we exposed below-saturation (*c*_0_ *< c*_*sat*_) concentrations of N-WASP solutions to supported lipid bilayers (1% Ni-NTA), and recorded the time series of N-WASP intensities. Figure 2c shows the fluorescent images at *t* = 10*min* and the corresponding kymographs of N-WASP intensities for 3 representative N-WASP solution concentrations (100 nM, 250 nM, 500 nM). At the beginning of experiments, we observed continuous association of N-WASP molecules onto the lipid bilayers. The mean pixel intensity appears to follow first-order surface binding kinetics, where the rate of surface association is proportional to the N-WASP bulk concentration (Fig. S7d, Fig. S8a-b). During this association process, the probability density histogram of pixel intensities shows a unimodal distribution (Fig. 2), consistent with N-WASP molecules individually adsorbing onto the lipid bilayer with homogeneous surface concentration. At a later time, N-WASP condensates appear on the lipid bilayer. Correspondingly, the probability density histogram of pixel intensities shows a bimodal distribution. This indicates two modes of surface association: individually adsorbed N-WASP molecules in the dilute phase (the lower-intensity peak in the histogram), and collectively-associated N-WASP molecules in the condensed phase (the high-intensity peak in the histogram; Fig. 2c). By analyzing this intensity histogram as a function of time, we find that the transition from unimodal to bimodal distribution takes the form of an abrupt switch: The peak representing the adsorbed phase decreases in amplitude, while a second peak rapidly emerges and shifts to higher intensities (Fig. 2e). These N-WASP condensates have layer indices that are consistently above 1 (Fig. S4d), suggesting that they are 3D surface objects in contact with the adsorbed layer. In addition, by photobleaching a region of lipid bilayers with associated N-WASP condensates, we verified that N-WASP molecules in the adsorbed layer exchange between the dilute and the condensed phase (Fig. S9). Together, this is consistent with the condensed phase of N-WASP on the surface forming via an abrupt switch-like transition from the adsorbed layer.

### Evidence for N-WASP surface condensates forming via a prewetting transition on the lipid bilayer

The theory of prewetting predicts that surface condensates form upon surpassing of a common critical surface concentration (31). Therefore, we set out to determine if such a critical concentration exists in our in vitro N-WASP condensation experiments. To obtain microscope images with an acceptable signal-to-noise ratio these were recorded under conditions that varied with the experimental condition, thus precluding a direct comparison of intensities. We developed two approaches to reliably determine the onset of condensation directly from imaging data. Firstly, we calculated the condensation rate as a function of time following N-WASP condensate segmentation, and extracted the time at which this rate peaks (Fig. 2d upper, Fig. S10a-b). Secondly, by fitting multivariate Gaussian functions to the time-dependent histogram of pixel intensities, we extracted the time at which N-WASP intensities no longer form a single peak characteristic of the dilute phase (Fig. 2e right, Fig. S11a-c, Methods). We find that with both approaches the time of condensation appears to scale inversely with the bulk concentrations, with a close to *−*1 exponent, 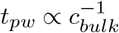 (Fig. 2f upper, Fig. S10d, Fig. S11d, Supplementary Information). This scaling coefficient is consistent with first-order adsorption kinetics, providing evidence that N-WASP condensation takes place around a common surface concentration. In addition, we find that the resultant normalized N-WASP fluorescence intensities, which are a measure of surface concentration, are of comparable values at this time of onset of condensation (Fig. 2f lower, Supplementary Information). Taken together, our data are consistent with a scenario where N-WASP condenses whenever the N-WASP surface concentration on the bilayer surpasses a critical value. This is indicative of N-WASP condensates forming on the lipid bilayers via prewetting transition.

### Actin polymerization counteracts the N-WASP prewetting transition

Actin nucleation via N-WASP and Arp2/3 is a major biochemical process for driving actin cortex formation (32, 33). We have observed both in vivo (20) and in vitro (Fig. 1e) that actin polymerization counteracts N-WASP condensation. We next ask if actin polymerization impacts on the N-WASP prewetting transition. We find that this effect is detectable when using a binary mixture of *C. elegans* (250 nM) and human N-WASP (250 nM, see SI). Both variants condense on the supported lipid bilayers and co-localize (Fig. S12). We compared the condensation kinetics of N-WASP only (Fig. 3a and Movie S6), versus the condensation kinetics of N-WASP in the presence of actin monomers and Arp2/3 complex (Fig. 3b and Movie S7). Similar to before (Fig. 2c) and as a hallmark of the prewetting transition, we observed in both conditions a split-up of the intensity histogram into a dilute and a condensed branch (Fig. 3c-d). In the presence of actin and Arp2/3, the time of histogram splitting is commensurate with the time of actin nucleation (Fig. S13). Interestingly, while the split is maintained without actin and Arp2/3, it is reversed at a later time in their presence (Fig. 3d, Fig. S13). This is indicative of an absence of N-WASP prewetting condensates at later times when actin filaments are present (see also Fig. S14-16, Movie S4 and Supplementary Information). Together, these results are consistent with actin polymerization, initiated in the condensates, suppressing the prewetting transition of N-WASP on lipid bilayers.

**Fig. 3.**
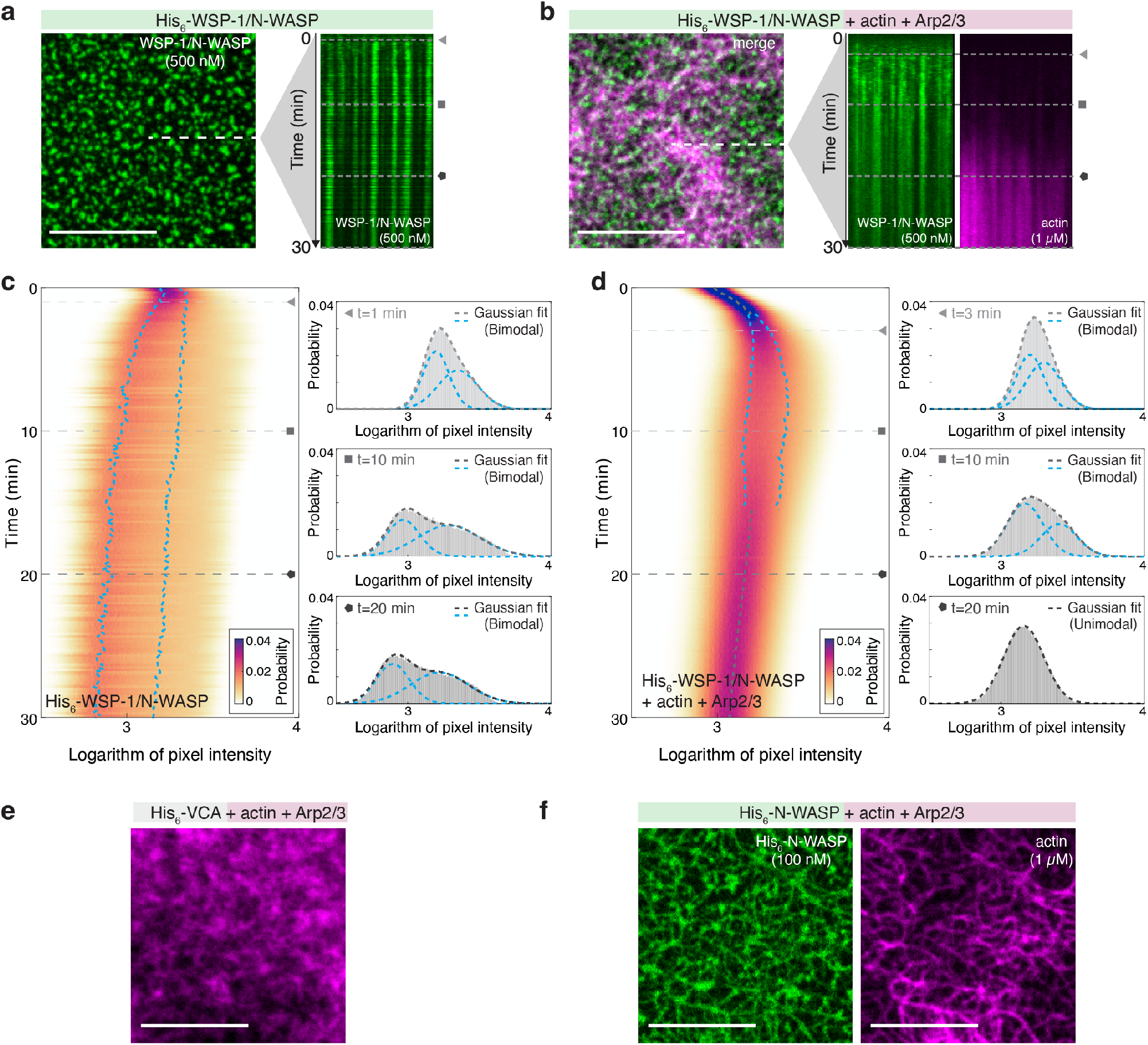
Actin polymerization counteracts prewetting. a) Confocal images of binary mixture of His6-WSP-1 and His6-N-WASP (500 nM, 10 % AF488 labeled, green) alone, and (b) in presence of actin (1 *μ*M, 10 % AF647 labeled, magenta) and Arp2/3 (10 nM) on SLBs after 30 min incubation. Kymographs along dotted line from 0 to 10 min (Movies S6 and S7, Figure S15). c) Left: Probability density kymographs of logarithm of N-WASP/WSP-1 pixel intensities for time-lapse images from a). Gray lines (single Gaussian) and blue lines (sum-of-two-Gaussians) represent overlays from either Gaussian-fitting method, determined for each time point by comparing the fitting residues between the two methods. Right: Histograms at 3 distinct time points (marked with triangle, square and circle) with either a single Gaussian or a sum-of-two-Gaussians fit (grey dashed lines). Blue dashed lines show respectively the fitted lower and higher Gaussian peaks in the case of a sum-of-two-Gaussians fits. d) Same as in c), Probability density kymographs of logarithm of N-WASP/WSP-1 pixel intensities for time-lapse images from the data with actin and Arp2/3 (b). e) Confocal images of actin network (1 *μ*M, 10 % AF647 labeled, magenta) formed for 30 min on SLB with Arp2/3 (10 nM) and 100 nM His6-VCA or f) full length His6-N-WASP (green). Scale bars 10 *μ*m.

### Altering cortex architecture

The actin cortex can display a complex architecture (34), and actin cortex structure is important for actin cortex functionality (35, 36). Can the nucleation of actin filaments from prewetting condensates alter the architecture of the resulting cortex, as compared to a homogeneously nucleated cortex? We formed in vitro cortices on supported lipid bilayers (1 % DGS-Ni-NTA) either in the presence of full length human N-WASP (100 nM) capable of forming prewetting condensates, or in the presence of the functional N-WASP domain VCA only (100 nM), which does not contain an intrisically disordered domain and does not form condensates (37). We find that the presence of VCA leads to the formation of dense actin meshworks on the bilayer (Fig. 3e). However, cortices formed in the presence of N-WASP appear to branch more sparsely, showing a significantly higher degree of bundling. They more closely resemble an in vivo actomyosin cortex (Fig. 3f) (38). We conclude that nucleating actin filaments from N-WASP prewetting condensates results in an altered cortex architecture.

## Discussion

Protein condensates can form inside cells at specific times and locations, to support e.g. specific chemical reactions (3, 39). How exactly the formation and dissolution of condensates is regulated remains unclear in many instances. Here we demonstrate that N-WASP at physiological concentrations can undergo a prewetting transition on lipid membranes, giving rise to N-WASP condensates that can drive actin polymerization reactions.

Prewetting transitions fall into the general class of wetting phenomena. Several examples of wetting phenomena have been observed in a biological context. Wetting of biological surfaces by protein condensates has been observed, for example, for transcription factors on DNA (28), for proteins on lipid membranes (40) and for proteins on microtubules (41). For the latter, TPX2 protein can wet microtubules and form a coat. Interestingly, this wetted layer has been shown to undergo a Rayleigh-Plateau instability, which drives the formation of evenly-spaced droplets (42). In wetting phenomena, both the surface affinity and interfacial tension plays a role in determining the condensate-surface contact morphology.

The prewetting transition, on the other hand, is driven solely by the affinity between protein molecules and biological surfaces. As such, the presence of surfaces promotes the appearance of the condensed phase at bulk concentration that are below the critical concentraion for bulk phase separation (28). In this paper, we report evidence that N-WASP undergoes a prewetting transition on lipid membranes, forming N-WASP condensates that are limited to the bilayer. These can drive the nucleation of actin filaments, giving rise to branched actin networks with a more bundled architecture as compared to networks that are formed in the absence of N-WASP surface condensation (similar to other phase-separation induced actin network structures (16, 17)). A key finding of the work here is that the growing actin meshwork itself appears to reverse the N-WASP prewetting transition. This is consistent with N-WASP proteins reentering the dilute phase, a key aspect of the dynamic instability of cortical condensates in vivo (20). We speculate that counteracting prewetting plays an important role in orchestrating the switch from growth to shrinkage of cortical condensates in vivo, which suppresses coarsening for collectively maintaining an emulsified state (20). Future work will be required to understand the precise molecular mechanisms that underlie this actin-driven suppression of N-WASP prewetting. One possibility is that as N-WASP molecules associate to Arp2/3 and drive actin filament nucleation, their ability to phase separate reduces due to changes of binding coefficients. A second possibility is that the actin filaments that grow from N-WASP surface condensates provide new surfaces for N-WASP association that compete with the lipid bilayers for N-WASP binding.

Taken together, we suggest that N-WASP prewetting and its suppression by actin is important for the shaping of a functional actomyosin cortex architecture. Given the observed kinetics of N-WASP and actin condensates in *C. elegans* oocytes (20), it will be of interest to explore the potential role of prewetting for other processes of condensate formation in vivo.

## Material and Methods

Additional details of experiments and analyses are described in the Supplementary Information, which includes Figures S1-S16, Movies S1-S7 and Table S1.

### Protein expression and purification

N-WASP proteins were expressed in Sf9 cells using the baculovirus system (43). Untagged *C. elegans* WSP1 (UniProt Q17795, plasmid TH1828) was purified following a protocol for N-WASP purification of Ho et al. (44) (Supporting Fig. 1a). MBP-His6-WSP1 (plasmid TH2035), His6-MBP-mGFP-WSP1 (plasmid TH1614) and MBP-His6-N-WASP (human, UniProt O00401, plasmid TH2098) were purified by His- and MBP-affinity columns and subsequent size-exclusion chromatography and stored in SEC buffer (50 mM HEPES (pH 7.4), 500 mM KCl, 5 % glycerol, 1 mM DTT) as detailed in the supplementary information. For MBP-His6-WSP1 and MBP-His6-N-WASP the MBP moiety was cleaved off right before an experiment with 5 % (v/v) GST-3C (in-house; 1 U/*μ*l) for at least 30 min at room temperature. For His6-MBP-mGFP-WSP1 the MBP moiety was cleaved off with 5 % (v/v) TEV protease (in-house; 1 U/*μ*l) (Supporting Fig. S1b). The samples were filtered with 0.1 *μ*m PVDF centrifugal filters (UFC30VV, Merck) and the concentration was measured using adsorption at 280 nm and 488 nm on a Nano-Photometer (Implen, NP80). Actin was purified from rabbit muscle acetone powder following published protocols (45) and detailed in the supplementary information. Arp2/3 (from porcine brain, #RP01P) and cdc42 (his-tagged, human constitutively active mutant Q61L, #C6101-A) were purchased from Cytoskeleton Inc.

### Supported lipid bilayers

Supported lipid bilayers (SLBs) were formed from small unilamellar vesicles (SUVs) composed of Synthetic 1-palmitoyl-2-oleoyl-glycero-3-phosphocholine (DOPC), 0 - 5 % 1,2-dioleoyl-sn-glycero-3-[(N-(5-amino-1-carboxypentyl)iminodiacetic acid)succinyl] (nickel salt, DGS-NTA-Ni), 0.1 % 1,2-dioleoyl-sn-glycero-3-phosphoethanolamine-N-[methoxy(polyethyleneglycol)-5000] (ammonium salt) (PEG5000 PE) and 0.01 %DiL (all Avanti Polar Lipids). Lipids were dissolved in chloroform, mixed, desiccated over night and resuspended in SLB buffer at a final concentration of 1 mg/ml. (50 mM TRIS pH 7.5, 150 mM KCl, 1 mM DTT). SUVs were formed by 10x freeze thaw cycles in liquid N2 and 37 °C water bath, respectively, and subsequent extrusion through a 100 nm filter. SUVs were stored at -70 °C and thawed and spun down at 17000 xg for 45 min on the day of the experiment. 96-well glass-bottomed plates (Greiner Bio one, CellView 655891) were washed with Hellmanex III (Hëlma Analytics) over night and rinsed under a stream of MilliQ H2O. Individual wells were washed with 6 M KOH for 2x 1 h, thoroughly rinsed with MilliQ H2O, and equilibrated with 35 *μ*l SLB buffer. 15 *μ*L SUVs were added per well and incubated for 5 min. SLBs were washed 5 times with SLB buffer to remove excess SUVs. Buffer was exchanged right before the experiment to assay buffer and integrity of SLBs was checked by FRAP in each well.

### Phase separation assays

Freshly MBP-cleaved and filtered N-WASP proteins were prediluted to appropriate concentrations in SEC-buffer and 5 *μ*l were mixed rapidly with 20 *μ*l of actin polymerization buffer to obtain final concentrations of 15 mM Hepes, pH 7.4, 150 mM KCl, 0.1 mM MgCl_2_ and 0.2 mM ATP in low-binding eppendorf tubes by flickering. For experiments in the presence of actin, 5 *μ*M N-WASP were mixed with actin polymerization buffer containing actin (3 *μ*M final concentration, 5 % Alexa-647-labeled) and Arp2/3 (100 nM final concentration) keeping the total volume and buffer conditions constant. As a control latrunculin A was added at a final concentration of 50 *μ*M.

### Generation of phase diagrams

To generate a phase diagram of the proteins we followed the method of Fritsch et al. (46). 20 *μ*l of protein solution of three concentrations well above the critical point were transferred by pipetting with a cut tip into the wells of an ultra-low attachment 384-well plates (CellCarrier 384 Ultra, Perkin Elmer, #6057800) in triplicates. The plate was spun down at 200xg for 10 min with a swinging-bucket rotor before transferring to the microscope. Confocal z-stacks of the whole well area were captured with a 20x air objective (Olympus U Plan XApo 20x (0.8 NA) Air) on a spinning disk confocal microscope (Olympus IXPLORE, IX83 inverted motorized stand with hardware autofocus and stage-top z-piezo). Analysis of the condensed volume versus the total volume was carried out with a script by A. Fritsch in Matlab (46). The critical concentration c_sat_ is estimated by the axes intersection of the volume ratio of condensed protein (V_in_) over total volume (V_total_) versus bulk protein concentration. For the phase diagram of surface condensates on SLBs the condensed area fraction (A_in_ over total area A_total_ in the field of view) was measured by segmentation of condensates in Ilastik (see below). The critical concentration c_sat_ for surface condensation is estimated by the axes intersection of the condensed area fraction with the bulk protein concentration.

### Time-lapse imaging of in vitro condensates

For time-lapse imaging of N-WASP condensation and actin polymerization, the buffer was equipped with a cocktail of triplet state quenchers following a protocol of Usaj et al. (47). Therefore 10 ml of reaction buffer were prepared and mixed with trolox (freshly dissolved at 100 mM in methanol), cyclooctatetraene (COT) and 4-Nitrobenzyl alcohol (NBA) (both from 200 mM stock in DMSO) to reach 2 mM final concentration of each component. To form trolox-quinone the solution was exposed to UV-light (254 nm, 10 min under a sterile hood) in a 10 cm petri dish on ice. Subsequently, solution was filtered (0.2 *μ*m) and degassed (for 1 hr in an desiccator on ice). As oxygen scavenger system pyranose oxidase (1.4 mg/ml final concentration, NATE-1718, creative enzymes), catalase (0.01 mg/ml final concentration, C40) and 0.8% glucose were freshly mixed from frozen stocks and added to the reaction buffer 1 min before the start of the experiment. For ATP regeneration phosphocreatinine (10 mM, P7936) and creatine phosphokinase (53 U/ml, C3755) were added to the reaction mixture. Finally actin (10 % AF647 labeled) and Arp2/3 were added to the buffer at the indicated concentrations and the mixture was added to the wells containing supported lipid bilayers. N-WASP was added into the wells by pipetting with a cut tip 5 times up and down for mixing and imaging was start right after.

### Image acquisition

Confocal images and time-lapse movies were captured with a 100x Oil objective (Nikon, Plan Apochromat, DIC, 1.45 NA) on a spinning disk confocal microscope (Andor, inverted motorized Nikon stand with hardware autofocus) with 488, 561 and 647 laser lines. Time-lapse imaging was carried out in a single plane close to the cover glass and z-stacks were recorded after 1 hr. Images were processed using Fiji/ImageJ 2.9.0. For size analysis of condensates in bulk, maximum projections were generated and condensates were segmented in Ilastik.

### FRAP experiments

FRAP experiments were carried out on droplets with a radius between 1.5 and 3.5 *μ*m 5 min - 1 h after droplet formation. Photo-bleaching was performed for individual droplets using a 488 nm laser for 10x 50 *μ*s over the full area of the droplet. Recovery of fluorescence intensity was recorded at a rate of 5 sec/frame for 5 min. Data analysis was performed following the published protocol (48) and script of Lars Hubatsch, available here: https://gitlab.pks.mpg.de/mesoscopic-physics-of-life/DropletFRAP.

### Condensate segmentation in time-lapse N-WASP images

Condensed fractions of N-WASP microscopy images were segmented using the Ilastik (49) software (version 1.4.0rc8, *simple segmentation*). This neural network-based model was first trained with 1-2 representative snapshots from the image stack, then used to process the entire stack and generate segmentation results. For the kinetic assay of N-WASP surface condensation, the representative snapshot was taken as the first frame when condensates can be visually identified. In order to compare condensed fraction, a common model was trained from all representative snapshots (100 nM, 250 nM, 500 nM bulk concentrations) and then used for time-lapse segmentation. For the FRAP assay of N-WASP condensates on lipid bilayers, the representative snapshots were taken as the frame before and subsequent to photobleaching. In order to reliably quantify FRAP intensity kinetics from dilute and condensed phases, tracking of individual condensates was performed and used to correct the segmentation results for frame-to-frame shifts (SI methods). These segmentation results were finally exported from Ilastik and processed with custom Matlab scripts.

### Determination of layer index for N-WASP surface condensates

The layer index quantification was developed as an indirect measurement to determine if the observed surface condensates is close to being a single layer or multilayered. This quantification is enabled by the knowledge that the microscopy z resolution is on the same scale or larger than the typical thickness *≈*<1 *μ*m of N-WASP condensates observed from z-view. Therefore, fluorescent images acquired near the lipid bilayer is a close measurement of the volumetric fluoresence of surface condensates projected per area. For single-layer condensates, this volumetric fluorescence, integrated within the segmented condensate, shall scale linearly with the integrated condensate area (*V* ∝ *A*^*α*^, *α* = 1). For spherical condensates with a fixed contact angle instead, the volumetric and area fluorescence scale in a power law relation (*V* ∝ *A*^*α*^, *α* = 1.5). Thus, the value of such exponent *α*, which we name as the layer index, serves as an indicator of condensate layers. To extract this index, pairs of integrated area and integrated intensity values were acquired from all segmented condensates, then a power law fit was performed (SI methods).

### Quantification of N-WASP prewetting time

To capture the abrupt appearance time *t*_*pw*_ of N-WASP condensates on lipid bilayers, two metrics were adopted: the condensate number increase per area quantified from the segmentation result, and the identification of condensed phase directly from N-WASP fluorescence intensities. In the first metric, the time of maximum condensate number increase per unit area was taken as the abrupt switch time. For the second metric, the logarithm of N-WASP fluorescence intensities (SI methods, *x*) was first binned to calculate the probability distribution *P* (*x, t*) (∑_*x*_*P* (*x, t*) = 1). A univariate and a two-variate (sum-of-two) Gaussian fit were then performed to generate the fitted probability density functions: *F*_1_(*x, t*) and *F*_2_(*x, t*) (∫*F*_*i*_(*x, t*)*dx* = 1). The residue between the fitted probability density function and the intensity data can be determined, *r*_*i*_(*t*) = ∑ [*F*_*i*_(*x, t*)Δ*x − P*_*i*_(*x, t*)]^2^, where Δ*x* is the bin width. We note that compared to the two-variate Gaussian fit, a univariate fit from the same data distribution is bound to generate a larger residue value. Thus, an abrupt increase in the residue difference between the fit functions, *r*_0_(*t*) = *r*_1_(*t*) *− r*_2_(*t*), captures the abrupt appearance of the condensed N-WASP phase. The time of abrupt increase *t*_*pw*_ was therefore set as the time when this residue difference *r*_0_(*t*) increases above a common threshold (set to 1e-4 for N-WASP 100nM, 250nM, 500nM time series comparison and 2e-5 for WSP-1/N-WASP mix with or without actin comparison). Both metrics were first extracted per acquired image (time step: 5-10 s) and then spline-fitted to finer time steps (1 s). Finally, to offset the experimental time delay between bulk solution addition and start of image acquisition, we fitted the starting (0-30s) kinetics of mean N-WASP fluorescent intensity to linear surface association dynamics. The time at which the mean intensity crossed zero was then extracted as the delay *t*_0_, and was used in correcting for the switch time 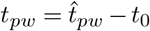 from both metrics.

### Quantification of N-WASP surface concentrations at prewetting

To convert N-WASP fluoresence intensities *I*_*s*_(*t*) into a measurement of surface concentration *c*_*s*_(*t*) = *ρ*_0_*I*_*s*_(*t*), a rescaling factor *ρ*_0_ is needed. To determine this rescaling factor, we note that the N-WASP surface association at low bulk concentrations closely resembles unsaturated first-order adsorption kinetics (SI methods). Thus, at the start of the surface association, N-WASP accumulation is approximately linear in time, *c*_*s*_(*t*) = *k*_*on*_*c*_*b*_*l · t*, where *c*_*b*_ is the applied bulk solution concentration and *l* the characteristic length for converting bulk to surface concentration units. *k*_*on*_, the association constant for N-WASP and lipid bilayers, stays a fixed number for datasets generated from the same experimental conditions (100 nM, 250 nM, 500 nM N-WASP bulk concentrations) and can thus be normalized. The fact that the exact values of *k*_*on*_, *k*_*off*_ (disassociation constant) and characteristic length *l* remain unknown hinder the rescaling of the mean N-WASP intensity to true surface concentration units. Yet, *ρ*_0_ can be determined such that for unsaturated adsorption (100 nM), the rescaled “surface concentration” *c*_*s*_(*t*) at long time is *c*_*b*_ (100 A.U.). An extended discussion on the unit conversion and interpretation is included in SI methods. Finally, the prewetting time *t*_*pw*_ determined previously from the N-WASP condensed phase appearance was used to extract the rescaled N-WASP surface concentration *ĉ*_*s*_(*t*_*pw*_) = *ρ*_0_*I*_*s*_(*t*_*pw*_) at prewetting.

## Data repository and material availability

Custom scripts are available at GitLab– The Open Research Data Repository of the Max Planck Society. Materials are available upon requests.

## ACKNOWLEDGEMENTS

S.W.G. was supported by the DFG (grant nos. TRR 83, GR 3271/2, GR 3271/3 and GR 3271/4) and the European Research Council (grant nos. 742712 and H2020-MSCA-ITN-2015). T.W. and J.L. thank the ELBE programm of MPI-PKS and MPI-CBG for funding and support. This work was funded by the Max Planck Society and received support from the DFG under Germany’s Excellence Strategy no. EXC-2068-390729961. We acknowledge the PEPC, LMF, HTS facility and the services of the MPI-CBG for their support. We thank A. Pozniakovsky, K. Crell, T. Neumann, V. Yan, A. Narayanan, D. Sun, L. Hubatsch, R. Alert, C. Weber, and F. Jülicher for experimental help and discussions. We are also grateful to A. Honigman and I. Seim for critical comments on the manuscript.

