## Supplementary Text and Figures for "Actin polymerization counteracts prewetting of N-WASP on supported lipid bilayers"

### Supplementary Note 1: Supplementary information on experimental methods

**A. N-WASP protein expression.** Sf9 cells (Expression Systems, 94-001F) infected with the respective baculovirus (1) were collected after 72 h by centrifugation for 30 min at 300 rpm. The cells were resuspended in lysis buffer (50 mM HEPES (pH 7.4), 500 mM KCl, 20 mM Imidazole, 5 % glycerol, 1 mM DTT (Dithiothreitol), 1 mM PMSF (phenylmethylsulfonyl fluoride), EDTA (ethylenediaminetetraacetic acid)-free protease inhibitor cocktail set III (Calbiochem) and 0.25 U/ml benzonase (in-house) and lysed 10x with a dounce homogenizer. The lysate was cleared by centrifugation for 30 min at 38,000 × g and 4 °C. The supernatant was filtered through 0.2 $\mu$m cellulose nitrate membranes (Whatman). MBP-His tagged proteins were run on a Ni-NTA column (Protino, Macherey-Nagel GmbH) at room temperature by a peristaltic pump. After washing with wash buffer I (50 mM HEPES (pH 7.4), 500 mM KCl, 20 mM Imidazole, 5 % glycerol, 1 mM DTT), his-tagged protein was eluted using His-elution buffer (50 mM HEPES (pH 7.4), 500 mM KCl, 250 mM Imidazole, 5% glycerol, 1 mM DTT). The eluate was further purified with amylose resin (NEB) in Econo-Pac gravity columns (Bio-Rad). After washing with His-elution buffer, MBP-tagged protein was eluted using MBP-elution buffer (50 mM HEPES (pH 7.4), 500 mM KCl, 250 mM Imidazole, 20 mM Maltose, 5 % glycerol, 1 mM DTT). The eluate was concentrated using Vivaspin 30,000 MWCO concentrators (GE Healthcare or Sartorius) and subjected to size-exclusion chromatography (SEC) at room temperature using a Superdex 200 increase column (GE Healthcare) and SEC buffer (50 mM HEPES (pH 7.4), 500 mM KCl, 5 % glycerol, 1 mM DTT). After concentrating the sample as described above, the proteins were stored at 4 °C for no longer than 2 weeks. MBP-His-WSP1 and MBP-His-N-WASP were buffer exchanged using Zeba Spin Desalting Columns (Thermo Scientific) into DTT free buffer for labeling with CF488A maleimide (Sigma) and Alexa647 C2 maleimide (Thermo Fischer), respectively, at equimolar ratio for 2 h at room temperature and exchanged back into SEC buffer the same way.

**B. Mix of different N-WASP variants.** Experimentally, the challenge in studying the effects of actin polymerization on N-WASP prewetting lies in having N-WASP prewetting and actin polymerization onset at a similar time span. To resolve this, we mixed the human and *C. elegans* N-WASP variants. This approach allowed us to tune the condensation kinetics, while delaying hardening of the surface associated condensates (experimental observation). Note that we observe a similar trend of actin suppressing the prewetting transition of human N-WASP only. However, the kinetics are slower and the transitions are thus less pronounced (Fig. S16).

**C. Actin purification.** 1 g of rabbit muscle acetone powder (Pel-freez Biologicals, 41995-2) was dissolved in rapidly stirring 24 mL of Ca-Buffer G (2 mM Tris-Cl, pH 8.0 at 25 °C, 0.2 mM ATP, 0.5 mM DTT, 1 mM NaAzide, 0.1 mM $CaCl_2$) and stirred at 0 °C for 30 min. The tissue was removed by centrifugation at 15000 rpm for 30 min at 4 °C and subsequent filtering of the supernatant through glass wool. Pellets were resuspend with a metal spatula in the original volume of Ca-Buffer G, stirred at 4 °C for 30 min and spun down again. The filtered supernatants were combined and stirred at 4 °C. 2.5 mL of 2 M KCl and 0.2 mL of 1 M $MgCl_2$ were added per 100 ml of supernatant to polymerize the actin at 4 °C for (final concentrations: 50 mM KCl and 2 mM $MgCl_2$). After 1 h 5.6 g KCl were added for every 100 mL of liquid (final concentrations 0.8 M KCl) to dissociate tropomyosin. After 30 min the solution was centrifuged in an ultracentrifuge at 100,000xg for 2 h at 2 °C to pellet actin filaments. The pellets were washed with 1 ml Ca-Buffer G, resuspend in 3-5 mL of Ca-Buffer G and homogenized with douncer 15-20 times. Actin was depolymerized by dialysis against 1 l of Ca-Buffer G for 3 days with 10,000 MWCO dialysis tubing (SnakeSkin, ThermoFischer). For labeleing, 100 $\mu$l of actin was buffer exchanged with DTT-free Ca-Buffer G in ZebaSpin Despalting Coloumns (7k MWCO, ThermFischer). Actin concentration was determinded at 290 nm in a Nanodrop and equimolar amounts of Alexa647-Maleimide or N-(1-Pyrene)Iodoacetamide (ThermoFischer) were incubated with actin for 1 h at RT. Subsequently, buffer was changed back to Ca-Buffer G and unbound dye was captured in ZebaSpin desalting coloumns. Monomeric actin was obtained in the early fractions of a size exclusion run on a Sephadex200 at 4 °C. Actin was stored at 4 °C and used within 2 weeks for polymerization assays or snapfrozen and stored at -80 °C for all other experiments. After thawing aggregates were removed by ultracentrifugation at 180 000xg for 1 h.

**D. Actin pyrene assays.** Actin pyrene assays have been carried out following McCall et al. (2). 5% actin pyrene solution was prepared in fresh CaBG and incubated for 1 h on ice followed by centrifugation ( 180 000xg, 1 h) to remove nucleation seeds. The concentration of the supernatant was determined in a Nanodrop and the solution was further diluted to 10 $\mu$M. Right before the assay Ca-ATP-actin is converted to Mg-ATP-actin by incubation with 10xME buffer for 2 min at RT and 30 $\mu$l were transferred per well in a 96 half-area well plate for final concentration of 2 $\mu$M. MgBG, KMEI, Arp2/3 (10 nM final concentration), cdc42 (250 nM final concentration, if applicable) and different variants of N-WASP protein (100 nM final concentration) were pre-mixed and added simultaneously with a multichannel pipette to the wells containing actin. The final reaction volume was 150 $\mu$l and final buffer matches 15 mM Hepes, pH 7.4, 150 mM KCl, 0.2 mM ATP, 0.1 mM $MgCl_2$. Actin assembly was monitored in a fluorescence plate reader (Tecan Spark 20M) via the timecourse of pyrene fluorescence (Ex: 340 +- 25 nm, Em: 405 +- 8 nm) with 15 s interval.

DRAFT

### Supplementary Note 2: Supplementary information on data analyses

**A. Fitting of N-WASP adsorption kinetics on lipid bilayers.** This section discusses the non-dimensional analyses of first-order membrane adsorption kinetics, which leads to our choice of fitting functions for the human N-WASP intensity data presented in Fig. 2. It also instructs the choice of rescaling method for retrieving the prewetting concentrations (next section).
To start with, we take the mean fluorescent intensity of N-WASP across all pixels in the field of view as a proxy for the concentration level of N-WASP proteins that is instantaneously associated with the lipid bilayer. Then, the kinetics of such mean intensity differs for when the surface association process resembles unsaturated or saturated binding kinetics.
We first perform a coarse estimation of the number of membrane binding sites versus the number of N-WASP molecules in the applied bulk solution. For all experiments, a volume of 60$\mu$l N-WASP solution is applied to a cylinder-shaped well where the bottom of the well (diameter: 6.49 mm) is covered by supported lipid bilayers. Using that lipid molecules have an average surface area of 70 Å$^2$ (3, 4) and that every 1 out of 100 lipid molecules can serve as binding sites for N-WASP (1% NTA SLB), we come to the estimation that for 100nM N-WASP bulk solution, a volume of 8 $\mu$l is needed for all lipid sites in SLB to be filled by N-WASP molecules. For 250nM and 500nM, the volume needed is 3.2 and 1.6 $\mu$l respectively. In other words, saturated binding shall dictate N-WASP mean intensity kinetics in all cases of bulk solution applications.
To analyze the features of saturated surface binding kinetics, we first investigate in the theory limit of unsaturated surface binding kinetics. The gain of surface-associated N-WASP molecules from the unsaturated binding of bulk N-WASP molecules onto the lipid sites follows the rate equation below:

$$\frac{dc_s}{dt} = k_{on}(c_b * l) - k_{off}c_s \tag{1}$$

where $c_s$ is the concentration of N-WASP molecules associated with the lipid surface (Unit: [nM]/$[\mu m]^2$) and $c_b$ the concentration in bulk solution (Unit: [nM]/$[\mu m]^3$). $l$ is a characteristic length that can be interpreted as the height of bulk solution layer in which N-WASP molecules can bind and unbind to the lipid surface with kinetic coefficients $k_{on}$ ($[s^{-1}]$) and $k_{off}$ ($[s^{-1}]$).
The nondimensional form of the rate equation follows:

$$\frac{dc_s^*}{dt^*} = 1 - \frac{k_{off}}{k_{on}}c_s^* \tag{2}$$

where $c_s^* = \frac{c_s}{c_b * l}$ and $t^* = t * k_{on}$ are the nondimensionalized surface concentration and time variables.
In the case of saturated binding kinetics, the rate equation and its nondimensionalized form can be written as: (Langmuir assumptions (5)))

$$\frac{dc_s}{dt} = \frac{c_{s0} - c_s}{c_{s0}}k_{on}(c_b * l) - k_{off}c_s \tag{3}$$

$$\frac{dc_s^*}{dt^*} = (1 - \frac{c_s^*}{c_{s0}^*}) - \frac{k_{off}}{k_{on}}c_s^* \tag{4}$$

and nondimensionalized variables $c_s^* = \frac{c_s}{c_b * l}$, $c_{s0}^* = \frac{c_{s0}}{c_b * l}$, $t^* = t * k_{on}$.
Notably, equation [4] has the same form of solution for the dynamics of $c_s^*$ compared to the equation for unsaturated binding kinetics [2], only effectively altering the disassociation coefficient $k_{off}$: $k_{off}^{'} = \frac{1}{c_{s0}^*} + \frac{k_{off}}{k_{on}}$. The solution to both non-dimensional N-WASP surface concentrations follows:

$$c_s^* = \frac{k_{on}}{k_{off}}(1 - e^{-\frac{k_{off}}{k_{on}}t}) \tag{5}$$

An example graph showing kinetics of this solution is presented in Fig. S8A. While the saturated and unsaturated surface concentration kinetics differ in their final values at equilibrium, the rate of association at beginning of the reactions is unanimously 1 for non-dimensional concentration $c_s^*$. In other words, the initial N-WASP surface-association rates across the application of 100nM, 250nM and 500nM (saturated) bulk N-WASP solutions are predicted to be a constant $\frac{dc_s}{dt}|_{t=0} = c_b|_{t=0} * l$ that is linearly dependent on the applied bulk solution concentrations $c_b|_{t=0}$.
Therefore, to fit the adsorption kinetics of mean N-WASP intensity data $I_s(t)$ from the experimental application of N-WASP bulk solutions, we use the mean intensity values at start of the time series ($t < 30s$) to extract a linear surface association rate $\frac{dI_s(t)}{dt}|_{t=0}$. As experimentally, a variable time delay $t_0$ exists between the start of bulk solution application and the start of time-lapse image acquisition, an intercept on time axis is also extracted accompanying this linear fit: $I_s(t) = \frac{dI_s(t)}{dt}|_{t=0}(t + t_0)$.
Fig. S7D shows the linear adsorption fit performed for N-WASP mean intensities, while Fig. S8B shows the rescaled surface intensity kinetics ($I_s^*(t)/(c_b(t)|_{t=0})$) where the extracted $t_0$ is offset and the initial rates of mean intensity increase per bulk concentration are normalized to 1 ($I_s(t)$ is converted to $I_s^*(t)$ after this normalization).

DRAFT

**B. Retrieval of N-WASP concentrations at prewetting transition.** To reliably retrieve the surface N-WASP concentration $c_{pw}$ at the time of observed abrupt switch in N-WASP pixel intensity distributions (the predicted critical point for prewetting transition), two quantities need to be sequentially determined: The time at which the abrupt switch takes place ($t_{pw}$), and the rescaling factor $\rho_0$ to convert N-WASP mean intensities into surface concentrations ($c_s(t) = \rho_0 \cdot I_s(t)$) at this abrupt switch time. The seeked critical surface concentration is then a combination of the two quantities: $c_{pw} = c_s(t)|_{t=t_{pw}} = \rho_0 \cdot I_s(t)|_{t=t_{pw}}$. We determined $t_{pw}$ and $\rho_0$ separately from two independent analyses of the N-WASP intensity data. $t_{pw}$ is extracted from comparing the residue differences between single Gaussian and sum-of-two-Gaussians fits of the histogram of pixel intensities (see Methods) and combining with the offset $t_0$ (previous section). This section discusses logistics for the extraction of the conversion factor $\rho_0$ and interpretation of the rescaled surface concentration unit.

The major twist for determining $\rho_0$ is that the laser power and z-focus positions used for capturing N-WASP presence near lipid bilayers are chosen to optimise the signal-to-noise ratio performance per experiment. For different N-WASP bulk concentrations, the surface N-WASP condensates that grow on SLB at later times have very different levels of pixel fluorescent intensities, thus such imaging setups are not held to fixed parameters across experiments. In addition, the exact values of association ($k_{on}$) and disassociation ($k_{off}$) constants for between N-WASP and lipid bilayers remain unknown in our system.

Yet, it is possible to circumvent such limits by integrating the analysis with appropriate assumptions derived from the adsorption theory (previous section). To begin with, we note that in the normalization of mean N-WASP surface intensity per bulk concentration (Fig. S8B), the intensity data is timed a conversion factor $\rho_s$ to make the slope 1 ($I_s^*(t) = \rho_s \cdot I_s(t)$). Using a shortened annotation $c_{b0} = c_b(t)|_{t=0}$ as the bulk concentration at the start of adsorption, this translates to:

$$\frac{dI_s^*(\hat{t})/c_{b0}}{d\hat{t}} = \rho_s \cdot \frac{dI_s(\hat{t})/c_{b0}}{d\hat{t}} = \frac{\rho_s}{\rho_0} \cdot \frac{dc_s(\hat{t})/c_{b0}}{d\hat{t}} = 1 \tag{6}$$

at start of the adsorption kinetics, where $\hat{t}$ is the offsetted experimental time, $\hat{t} = t - t_0$.

Comparing the kinetic solutions from equation [6] and the nondimensional theory prediction [2] side by side, we obtain the expression for the seeked mean-intensity-to-surface-concentration conversion factor $\rho_0$:

$$\rho_0 = \frac{\rho_s}{c_{b0}} \cdot \frac{t^*}{\hat{t}} \cdot \frac{c_s(\hat{t})}{c_s^*(t^*)} = \frac{\rho_s}{c_{b0}} \cdot k_{on} \cdot (c_{b0} \cdot l) = \rho_s \cdot k_{on} \cdot l \tag{7}$$

$\rho_s$ can be directly determined from each N-WASP intensity time series as a normalization factor ($\rho_s = c_{b0}/(\frac{dI_s(t)}{dt}|_{t=0})$). For $k_{on}$ and the characteristic length $l$, they are parameters describing the surface affinity between N-WASP molecules and lipid bilayers for the in vitro assay, and thus can be reasonably assumed to stay constant across our experimental cases of applying different bulk N-WASP concentrations. In other words, the choice of their values do not interfere with the comparison of $\rho_0$ values, and thus the rescaled N-WASP surface concentrations, across datasets.

Nevertheless, an intuitive choice of $k_{on} \cdot l$ value comes from the recognition of binding kinetics in the case of applying 100nM bulk N-WASP solution. As predicted from equation [5], the final surface concentration in this case, $c(\hat{t})|_{\hat{t}\to\infty}$, is a well-defined quantity $c(\hat{t})|_{\hat{t}\to\infty} = \frac{k_{on}}{k_{off}} \cdot (100[nM]) \cdot l$. By choosing a value for the $k_{on} \cdot l$ such that the final N-WASP mean intensity $I(\hat{t})|_{\hat{t}\to\infty}$ is 100 A.U. for 100nM N-WASP bulk solution, we effectively rescale the equilibrium constant $(k_{on}/k_{off}) \cdot l$ to be 1 in the converted concentration unit. This choice of value $k_{on} \cdot l$ is then fixed and combined with the dataset-specific parameter $\rho_s$ to calculate the rescaled surface concentrations for all bulk N-WASP concentrations (100nM, 250nM, 500nM) as shown in Fig. 2f (lower).

**C. Coefficient of variation analysis and cross-correlation of N-WASP and actin fluorescence intensities.** To follow the kinetics of N-WASP condensation and actin polymerization we analysed the coefficient of variation (CV), defined as standard deviation devided by mean of pixel intensities in a field of view (Fig. S14c).

To evaluate the colocalization of N-WASP and actin as a function of time, we calculated the cross-correlation coefficient between the normalized N-WASP and actin fields at every time point. The normalization of the fields was performed by extracting the mean and dividing by the standard deviation from all pixels in the field of view (Fig. S16a). The cross-correlation coefficient was then calculated as the mean of the cross product between normalized N-WASP and normalized actin fields across all pixels in the field of view (Fig. S16b).

DRAFT

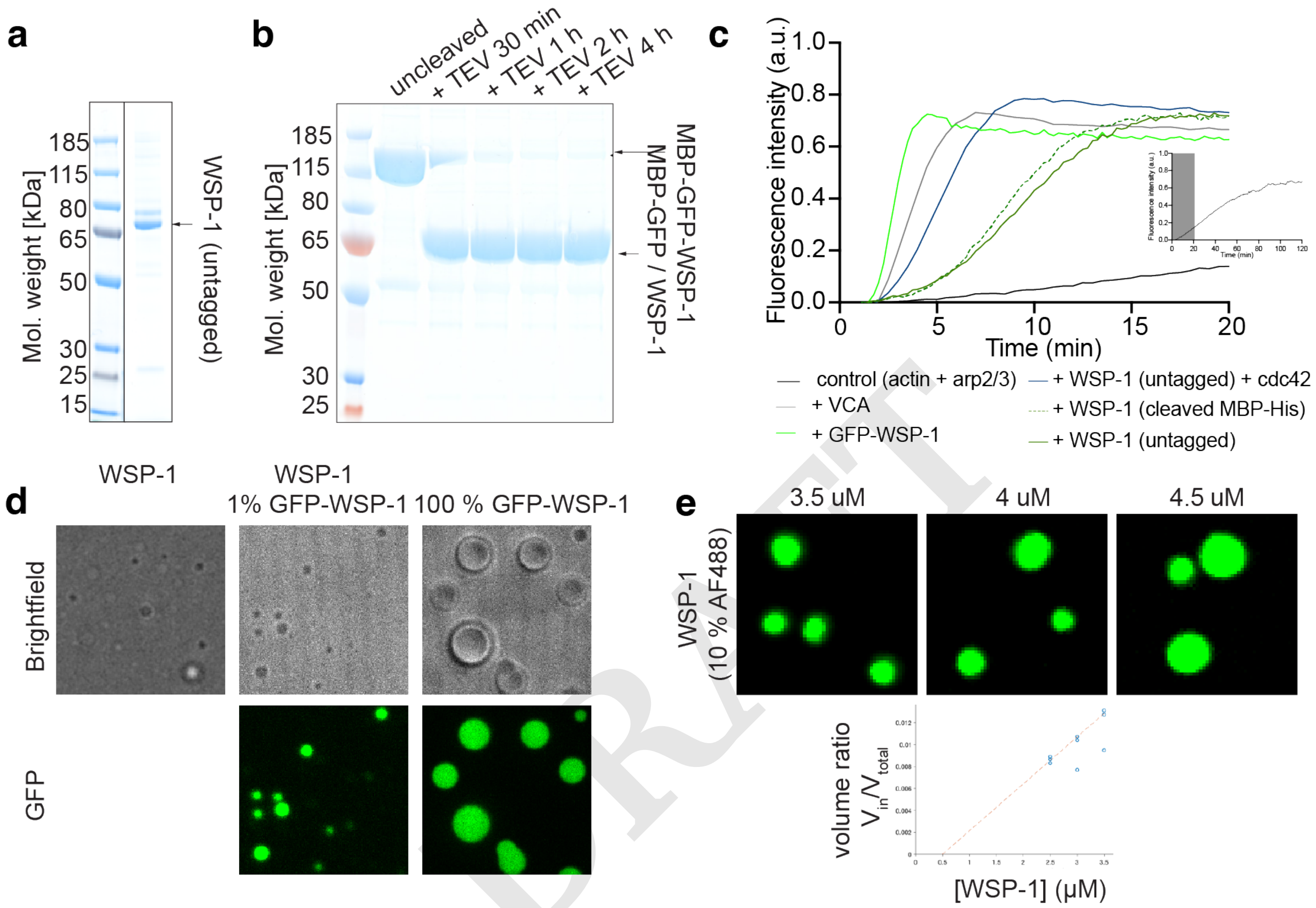


**Fig. S1. Characterization of recombinant *C. elegans* N-WASP (WSP-1).** a) SDS-Page of WSP-1 without additional tags, recombinantly expressed and purified from SF9 cells. b) SDS-Page of WSP-1 with N-terminal MBP-GFP tag, that was cleaved for 30 min, 1, 2 or 4 h by TEV protease at room temperature. c) Actin-pyrene assay of differently purified WSP-1 variants. Actin (2 $\mu$M) + Arp2/3 (10 nM) shows slow polymerization (black curve and insert). Addition of VCA domain (100 nM, gray) increases dynamics. Full-length WSP-1 (100 nM each, untagged in dark green, freshly cleaved from MBP-His$_6$-tag in dotted green) shows basal activity. Addition of cdc42 (constructively active mutant Q61L, 250 nM, blue) activates WSP-1 and increases actin polymerization kinetics. GFP-tagged WSP-1 (100 nM, light green) shows highest activity. d) Phase separation assay of untagged (left), mixture of untagged with 1% GFP-tagged (middle) and 100% GFP-tagged WSP-1 (each 5 $\mu$M in actin polymerization buffer). GFP-tagged WSP-1 shows increased phase separation propensity. To avoid the effects coming from the tag we used AF488-labeled N-WASP for subsequent experiments. e) Representative images and quantification of phase separation assay of His$_6$-WSP-1 labeled with 10 % AF488 (see Methods).

DRAFT

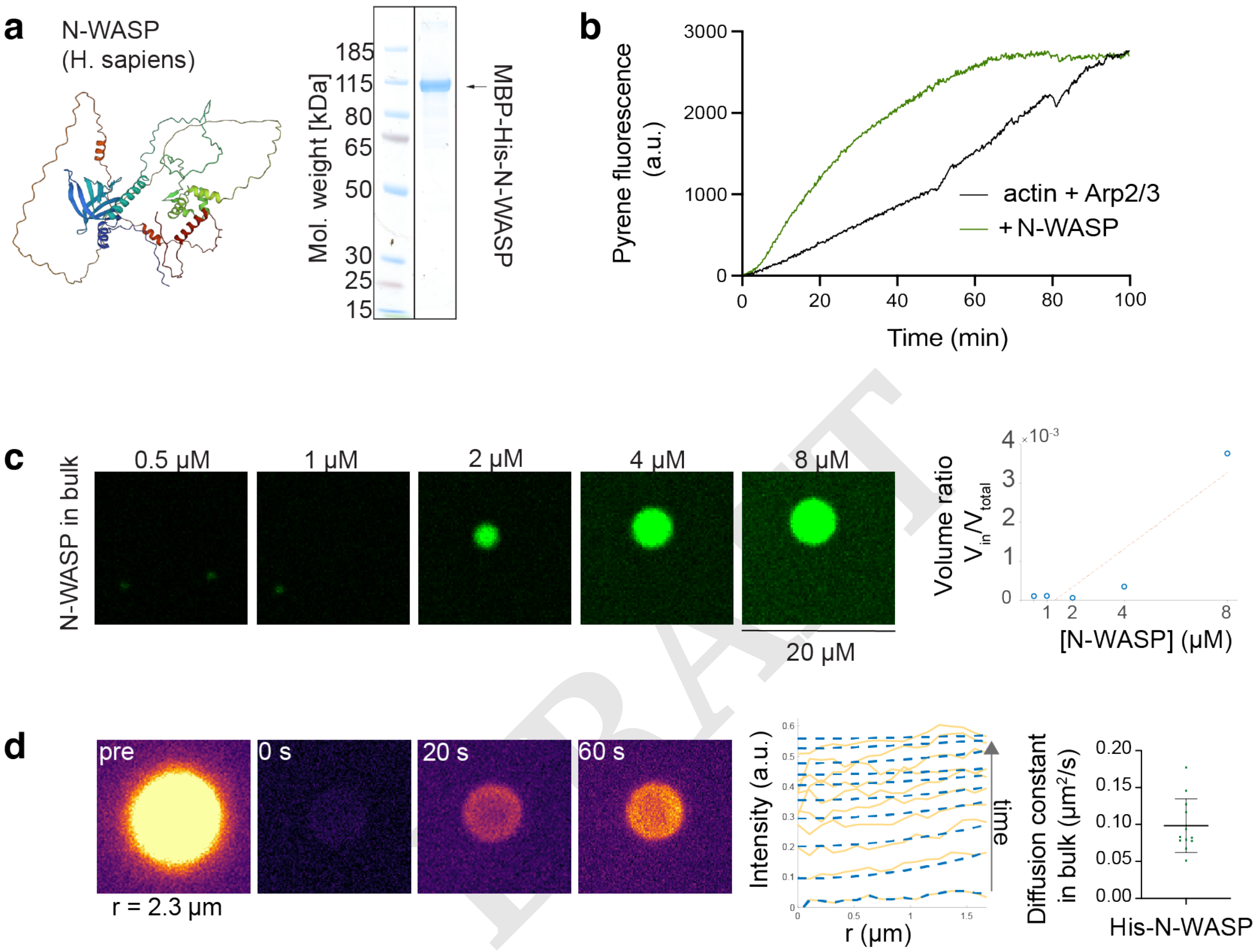


**Fig. S2. Characterization of recombinant *H. sapiens* N-WASP.** a) Alpha-fold prediction of human N-WASP with sequence-dependent color code. SDS-Page of MBP-$His_6$-tagged human N-WASP, recombinantly expressed and purified from SF9 cells. b) Actin-pyrene assay of actin (2 $\mu$M) + Arp2/3 (10 nM) alone (black curve) and in presence of full-length N-WASP (100 nM, freshly cleaved from MBP-$His_6$-tag, green curve), showing basal activity. c) Representative images and quantification of phase separation assay of $His_6$-N-WASP labeled with 10 % AF488 (see Methods). d) Left: Fluorescence recovery after photo-bleaching (FRAP) images of condensate formed from 5 $\mu$M $His_6$-N-WASP. Middle: Example plot of average fluorescence intensities (yellow lines) over the radius r in one condensate for 10 consecutive time points. The fits (blue dotted lines) were used for the model of protein diffusion in condensates to extract the diffusion coefficients (see Methods). Right: Diffusion coefficients D inside the condensates were determined for n = 12 condensates. Data are the mean ± s.d.

DRAFT

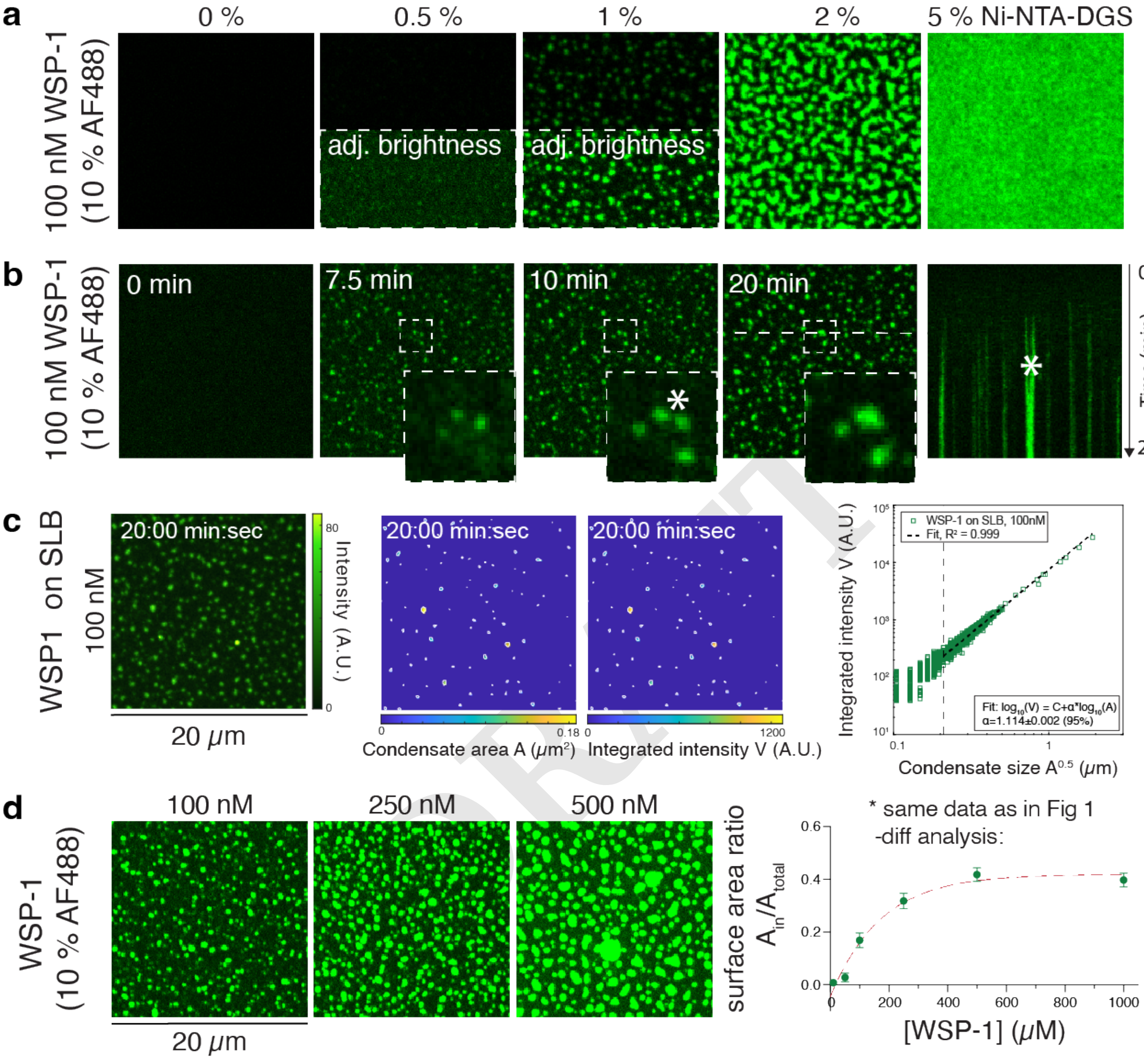


**Fig. S3. Detailed analysis of WSP-1 surface condensation on supported lipid bilayers.** a) Confocal images of the surface layer of $His_6$-WSP-1 (100 nM) interacting with SLBs doped with different concentrations of Ni-NTA-DGS (0, 0.5, 1, 2, 5 %) at steady-state (after 30 min). b) Confocal snapshots of a timelapse of the condensation of $His_6$-WSP-1 (100 nM) on SLB with 1 % Ni-NTA. Inset and asteriks pointing towards fusion event of 2 condensates on the membrane. Kymograph from 0 to 20 min along the dotted line. c) Confocal snapshot at t = 20 min, segmented condensates with color-coded mean area and mean integrated intensity, respectively. Plot of the integrated intensity compared to the respective condensate area. d) Confocal snapshots of $His_6$-WSP-1 condensates formed from different bulk concentrations (100, 250, 500 nM) on SLB with 1 % Ni-NTA at steady state (after 30 min). Plot of the surface area ratio of condensed ($A_{in}$) versus total Area ($A_{total}$) with exponential fit (red dotted line). Data are the mean ± s.d. All images are 20 $\mu$m in length.

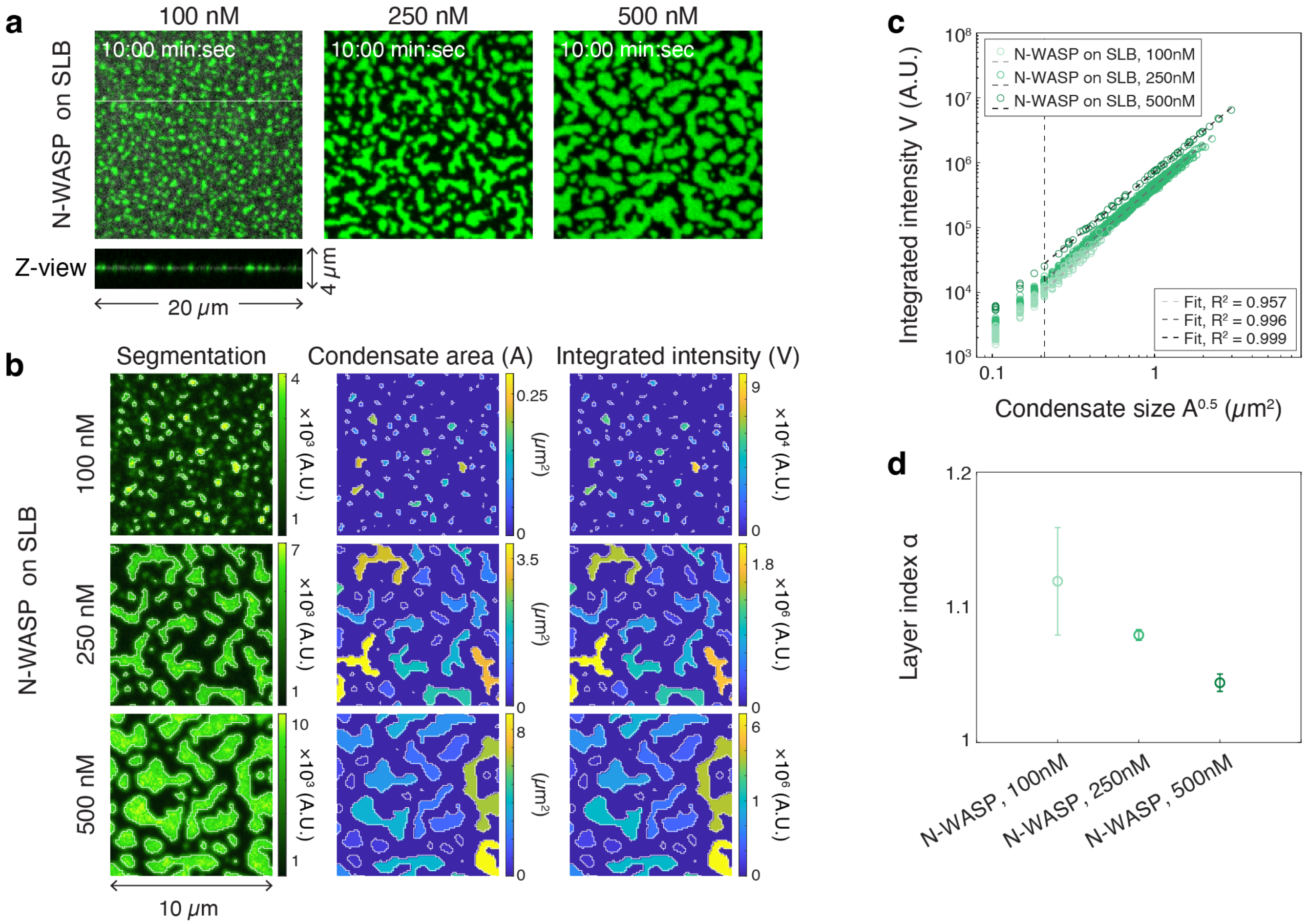


**Fig. S4. N-WASP forms multilayered surface condensates.** a) Confocal snapshots of a timelapse of the condensation of $His_6$-N-WASP at 100, 250 and 500 nM on SLB with 1 % Ni-NTA. b) Maximum projection and x-z-view of a confocal z-stack across the surface layer of 100 nM $His_6$-N-WASP condensates on a SLB with 1 % Ni-NTA. c) Confocal snapshots, segmented condensates with mean intensity, condensate mean area and integrated intensity for three different N-WASP concentrations. d) Plot of the Integrated intensity compared to the respective condensate area. e) Layer indices as revealed from d) for the three different N-WASP concentrations.

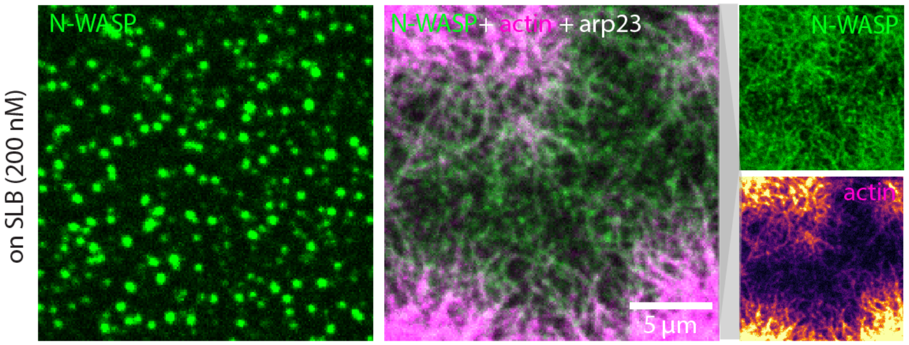


**Fig. S5. Actin controls shape of N-WASP surface condensates.** a) Confocal images of $His_6$-N-WASP (200 nM, 10% AF488 labeled, green) forming surface condensates on SLBs with 1 %Ni-NTA alone, or in presence of actin (1 $\mu$M, 10% AF647 labeled, magenta) and Arp2/3 (10 nM) at steady-state (after 30 min incubation).

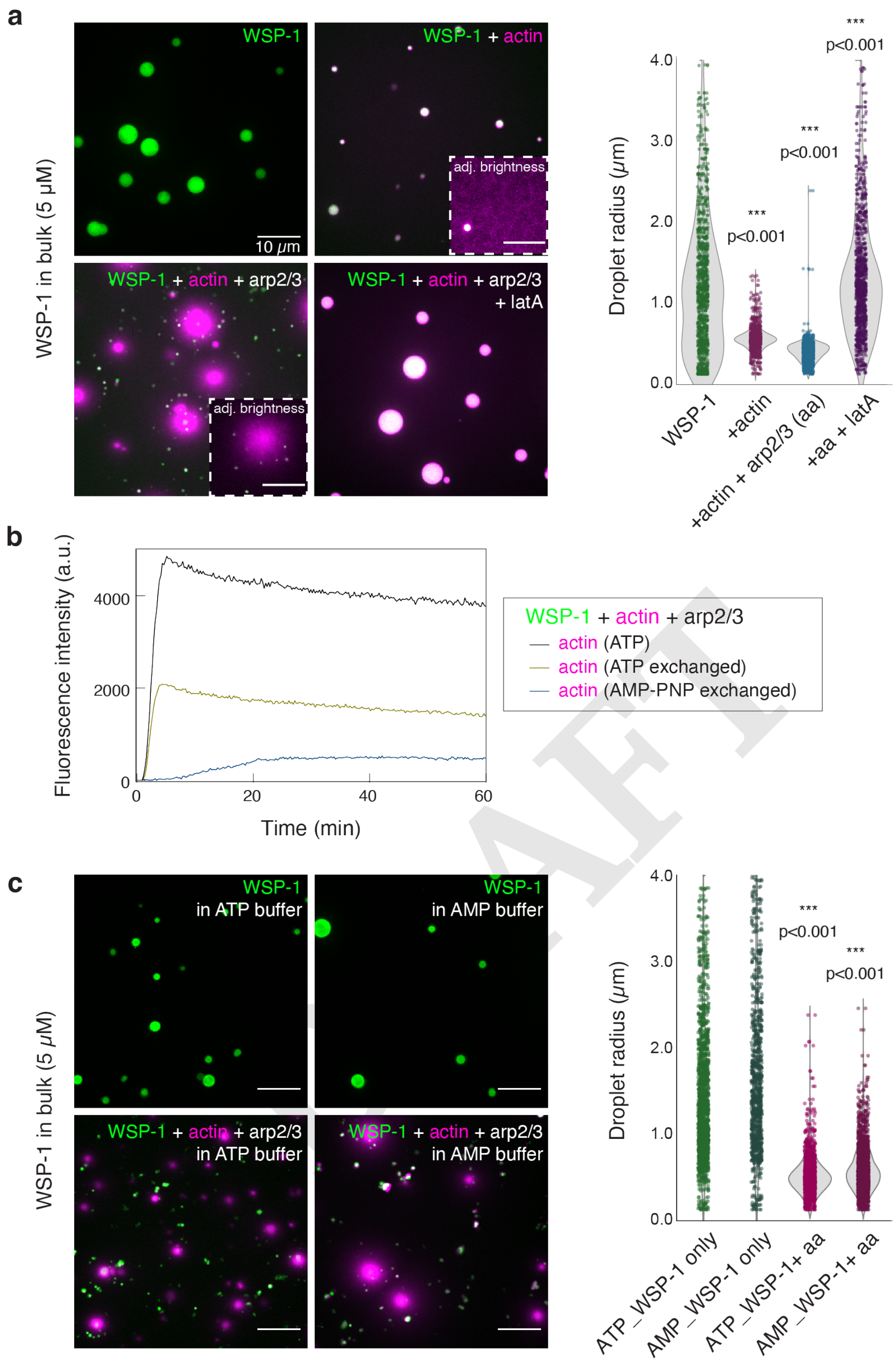


**Fig. S6. Actin polymerization controls size of WSP-1 condensates in bulk.** a) Left: Maximum intensity projections of confocal z-stacks of His$_6$-WSP-1 (5 $\mu$M, 10 %AF488 labeled) condensates formed over 1 h in bulk (upper row) in actin polymerization buffer alone (left images), and in the presence of actin (3 $\mu$M, 10 % AF647 labeled, magenta), actin and Arp2/3 (100 nM), and in the presence of latrunculinA (50 $\mu$M). Right: Quantification of droplet radii from segmented droplets with p-values obtained by one-way ANOVA analysis. b) Actin-pyrene assay in the presence of WSP-1 (100 nM) and Arp2/3 (10 nM). 2 $\mu$M of actin were either ATP-bound (black curve), nucleotide-exchanged with fresh ATP (yellow curve) or nucleotide-exchanged to non-hydrolysable AMP-PNP. c) Left: Maximum projections of confocal z-stacks of His$_6$-WSP-1 (5 $\mu$M, 10 %AF488 labeled) condensates formed over 1 h in bulk (upper row) in actin polymerization buffer with ATP (left images) or with AMP (right images), and in the presence of actin (3 $\mu$M, 10 % AF647 labeled, ATP bound versus AMP-PNP bound, magenta) and Arp2/3 (100 nM) (lower images). Right: Quantification of droplet radii from segmented droplets with p-values obtained by one-way ANOVA analysis. All scale bars 10 $\mu$m.

DRAFT

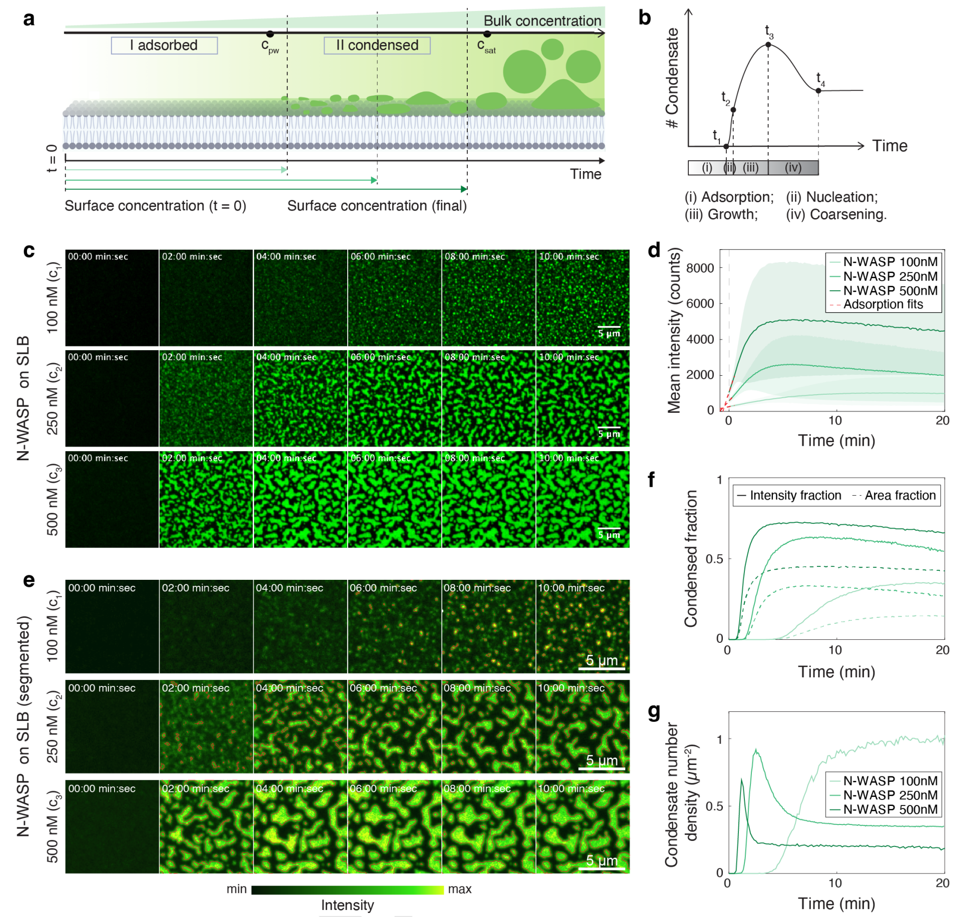


**Fig. S7. Quantification of N-WASP condensation dynamics on SLBs.** a) In the kinetic assay, N-WASP concentration near surface accumulates from zero at $t = 0$. The SLB surface goes through the adsorbed phase before condensation starts. b) Four stages is predicted for the kinetic assay of N-WASP surface condensation: Adsorption, nucleation, growth and coarsening. c) Time-lapse surface snapshots at different bulk concentrations of N-WASP (100, 250, 500 nM). Scale bar = 5 $\mu$m. d) The first 30s of the fluorescence intensity kinetics representing N-WASP surface association can be fitted to linear adsorption kinetics (red dashed lines). The offset time $t_0$ is used for subsequent correction of nucleation and split times. Shaded area represents standard deviation across intensities for all pixels in the field of view. e) Segmentation of surface condensates (red lines) in time-lapse snapshots. A same trained model (ilastik) is used to generate segmentation of N-WASP images with different bulk concentrations. Scale bar = 5 $\mu$m. f) Intensity and area fraction of segmented condensates (e) as a function of time. g) Area-averaged number density of segmented condensates (e) as a function of time.

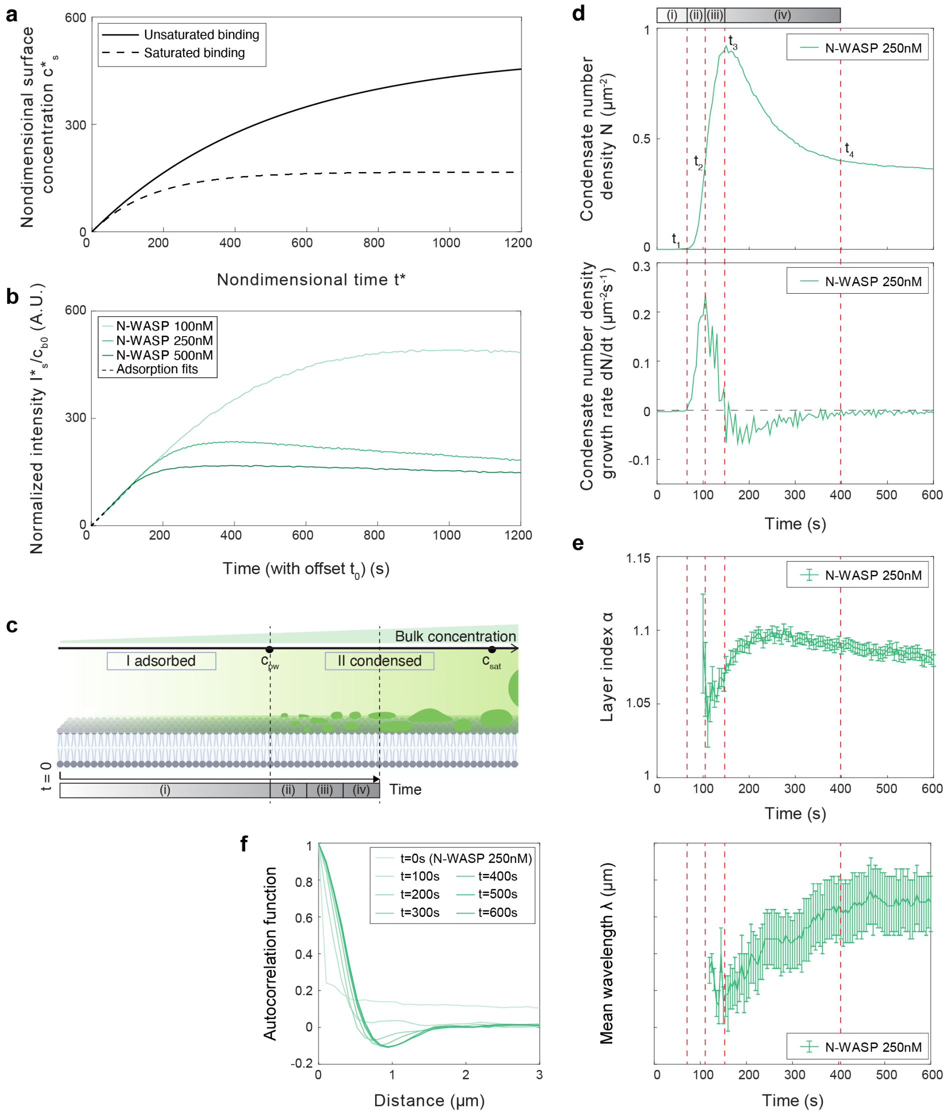


**Fig. S8. Quantification of N-WASP adsorption and condensation stages on SLBs.** a) Nondimensionalized protein surface concentration $c_s^*$ as a function of nondimensionalized time $t^*$ undergoing unsaturated (solid) or saturated (dashed) binding kinetics. $k_{on}/k_{off}$ is taken to be 500 for both scenarios. For the saturated binding scenario, an additional nondimensional saturation coefficient $c_0^* = 250$ is applied. b) Kinetics of mean N-WASP intensities normalized first by bulk concentration and then by enforcing the fitted adsorption rate to have a slope of 1. c) For individual condensates, the four stages of surface condensation map to dominance of different dynamics: (i) adsorption, no condensate appearance; (ii) nucleation, peaking individual condensate appearance rate; (iii) growth, individual condensate grows in size and height; (iv) coarsening, nearby condensates merge, mean size of condensate increases while total condensate number decreases. d) Take 250 nM N-WASP bulk concentration as example, start and end of the four stages $(t_1, t_2, t_3, t_4)$ can be extracted from the kinetics of condensate number (upper) and condensate number increase rate (lower). e) Layer index $\alpha$ extracted from (c) for segmented condensates as a function of time. f) Left: Spatially-averaged pixel-pixel intensity correlation plot for time-lapse fluorescence images. Right: Mean wavelength extracted from (f) left as a function of time.

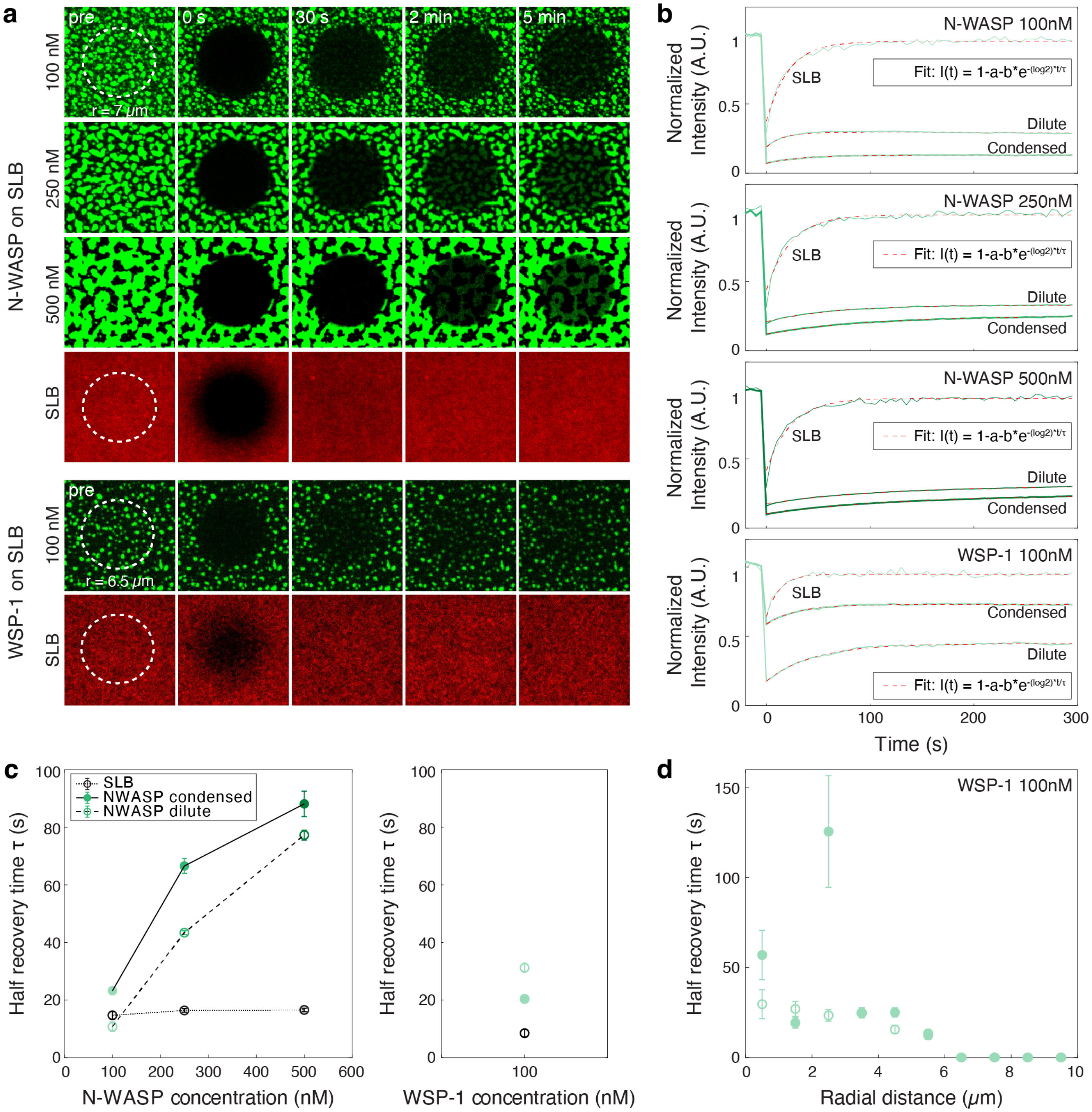


**Fig. S9. Quantification of N-WASP diffusion in and outside surface condensates.** a) Confocal snapshots of a FRAP experiment on SLBs with 1 %Ni-NTA, DiI and different concentrations of N-WASP (100, 250, 500 nM) and WSP-1 (100 nM). b) Fluorescence intensities over time of the DiI incorporated into the SLBs and the N-WASP signal in the dilute and condensed phase, respectively. c) Recovery half times extracted from the fits in b). d) Recovery times of the individual WSP-1 (100 nM) surface condensates with respect to their distance from the center of the bleaching spot.

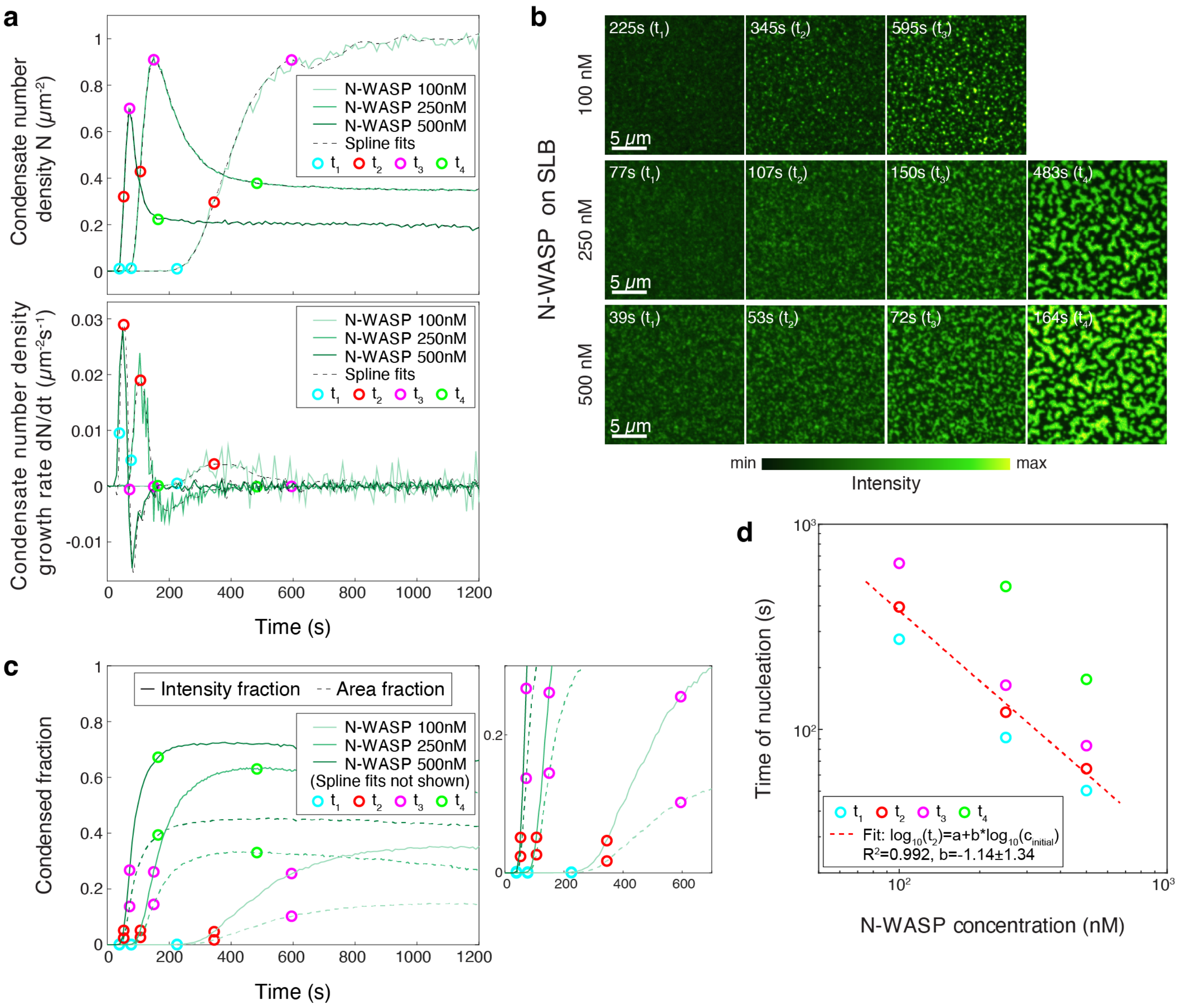


**Fig. S10. Segmentation-dependent quantification of critical N-WASP condensation on SLBs.** a) Upper: Condensate number density as a function of time for different N-WASP bulk concentrations (100, 250, 500 nM, same as S7g) overlayed with extracted stage times $t_1 - t_4$ (coloured circles) from spline fits (dashed lines). Lower: Condensate number increase rate as a function of time overlayed with extracted stage times. b) Snapshot comparison at $t_1 - t_4$ across different N-WASP bulk concentrations. Scale bar = 5 $\mu$m. c) Intensity and area fraction of segmented condensates as a function of time (same as S7f) overlayed with extracted stage times $t_1 - t_4$. Zoom-in plot (right) shows the comparable condensed fractions across different N-WASP bulk concentrations. d) Stage times $t_1 - t_4$ plotted as a function of N-WASP bulk concentrations. The nucleation time $t_2$ can be fitted to an inverse scaling relation (red dashed line).

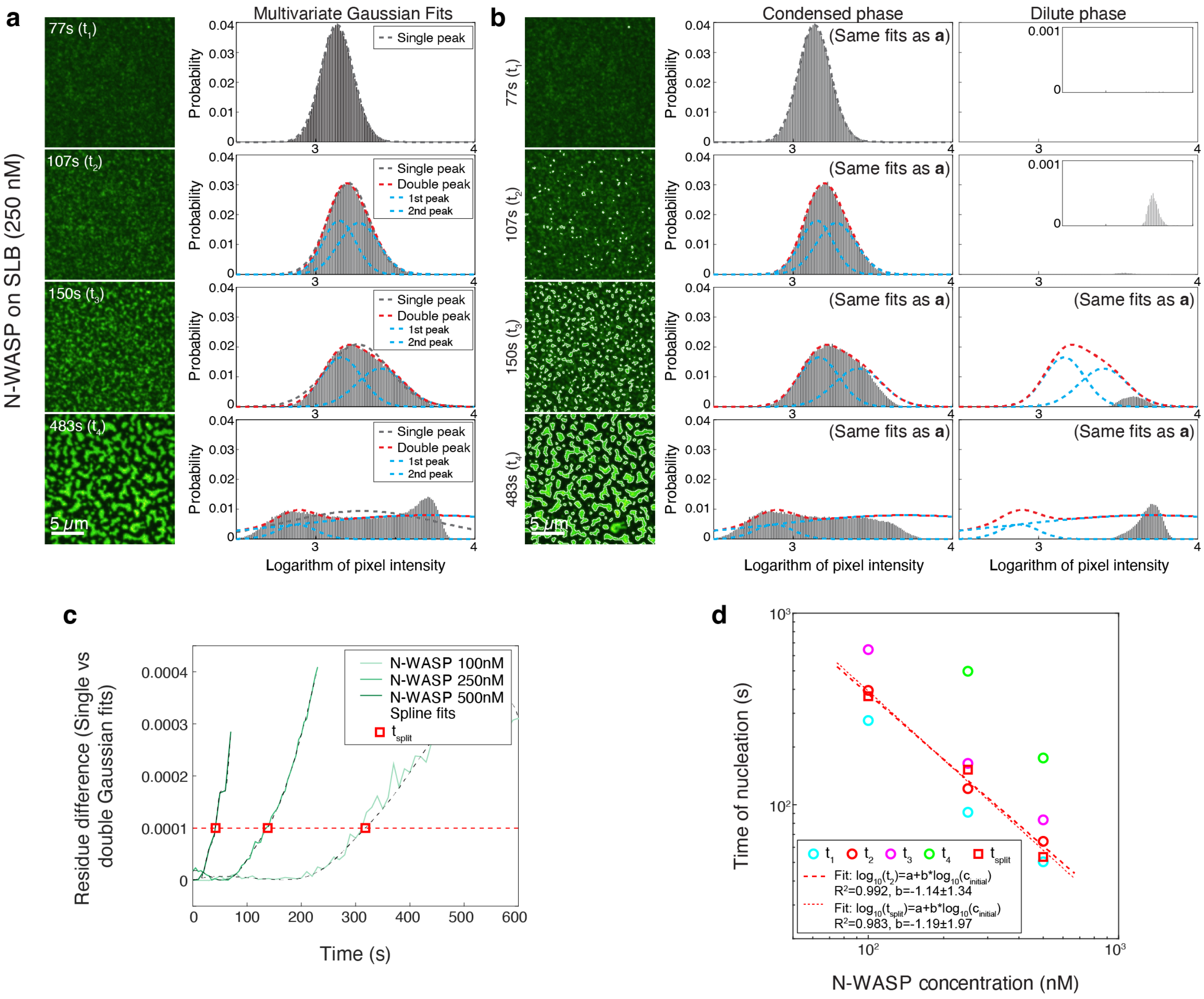


**Fig. S11. Segmentation-independent quantification of critical N-WASP condensation on SLBs.** a) Take 250 nM N-WASP bulk concentration as example, single-variate or double-variate Gaussian functions fits for the histograms of pixel intensity at different stage times $t_1 - t_4$. Scale bar = 5 $\mu$m. b) Comparison of fit from (a) with histograms of pixel intensity for segmented condensed and dilute phases at different stage times $t_1 - t_4$. Scale bar = 5 $\mu$m. c) Residue difference between single-variate and double-variate Gaussian fits (a) as a function of time. The residue difference can be used to extract the split time $t_{split}$ (red squares) after setting a threshold (red dashed lines). d) Comparison of split time $t_{split}$ and nucleation time $t_2$ plotted as a function of N-WASP bulk concentrations. Similar to S10d, the split time $t_{split}$ can also be fitted to an inverse scaling relation (red dotted line).

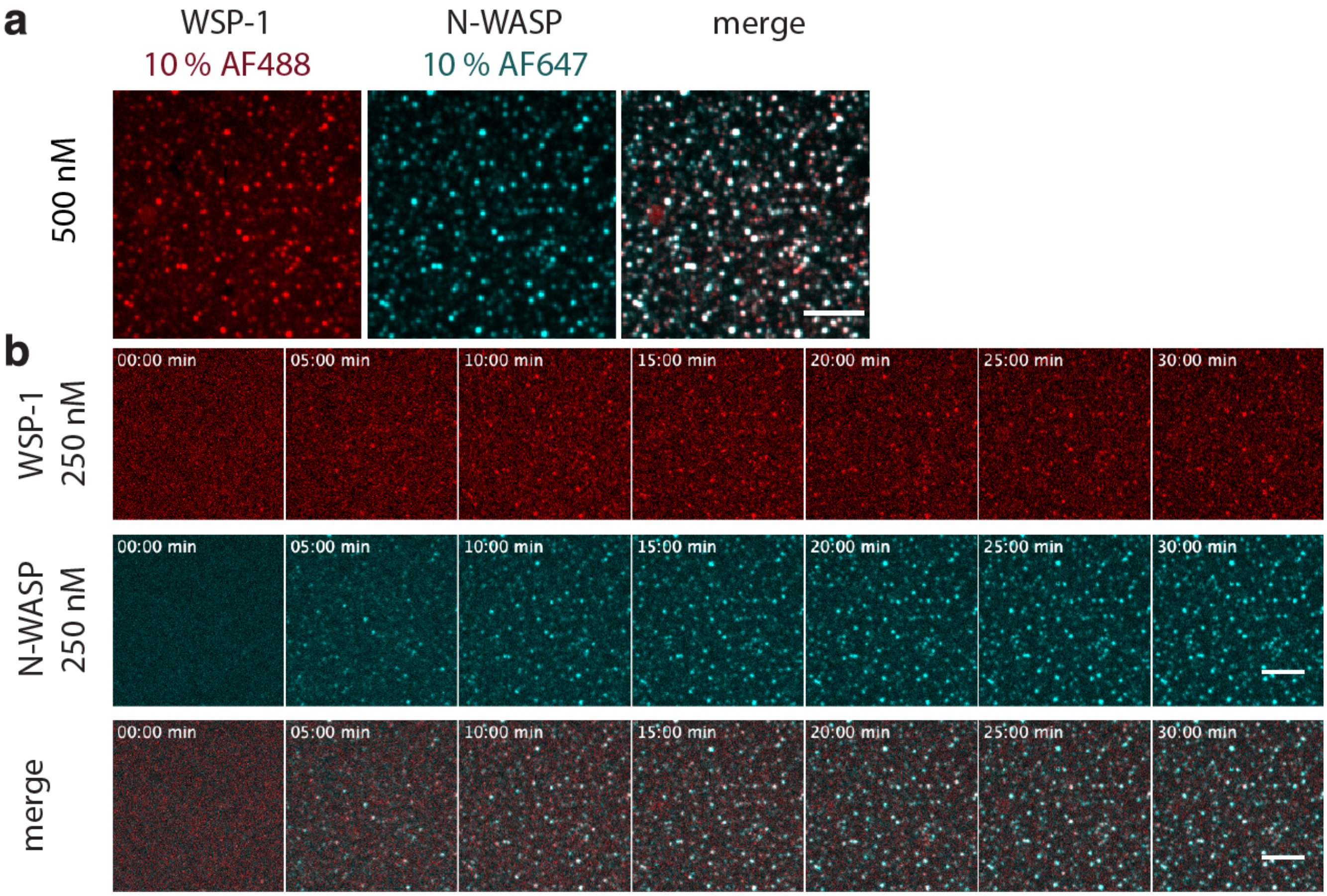


**Fig. S12. Human and *C. elegans* N-WASP form surface co-condensates.** a) Confocal images of *C. elegans* WSP-1 (250 nM, 10 %AF488 labeled, red) and *H. sapiens* N-WASP (250 nM, 10 %AF647 labeled, turquoise) b) Confocal snapshots of a timelapse of the condensation of $His_6$-N-WASP at 100, 250 and 500 nM on SLB with 1 % Ni-NTA. All scale bars = 10 $\mu$m.

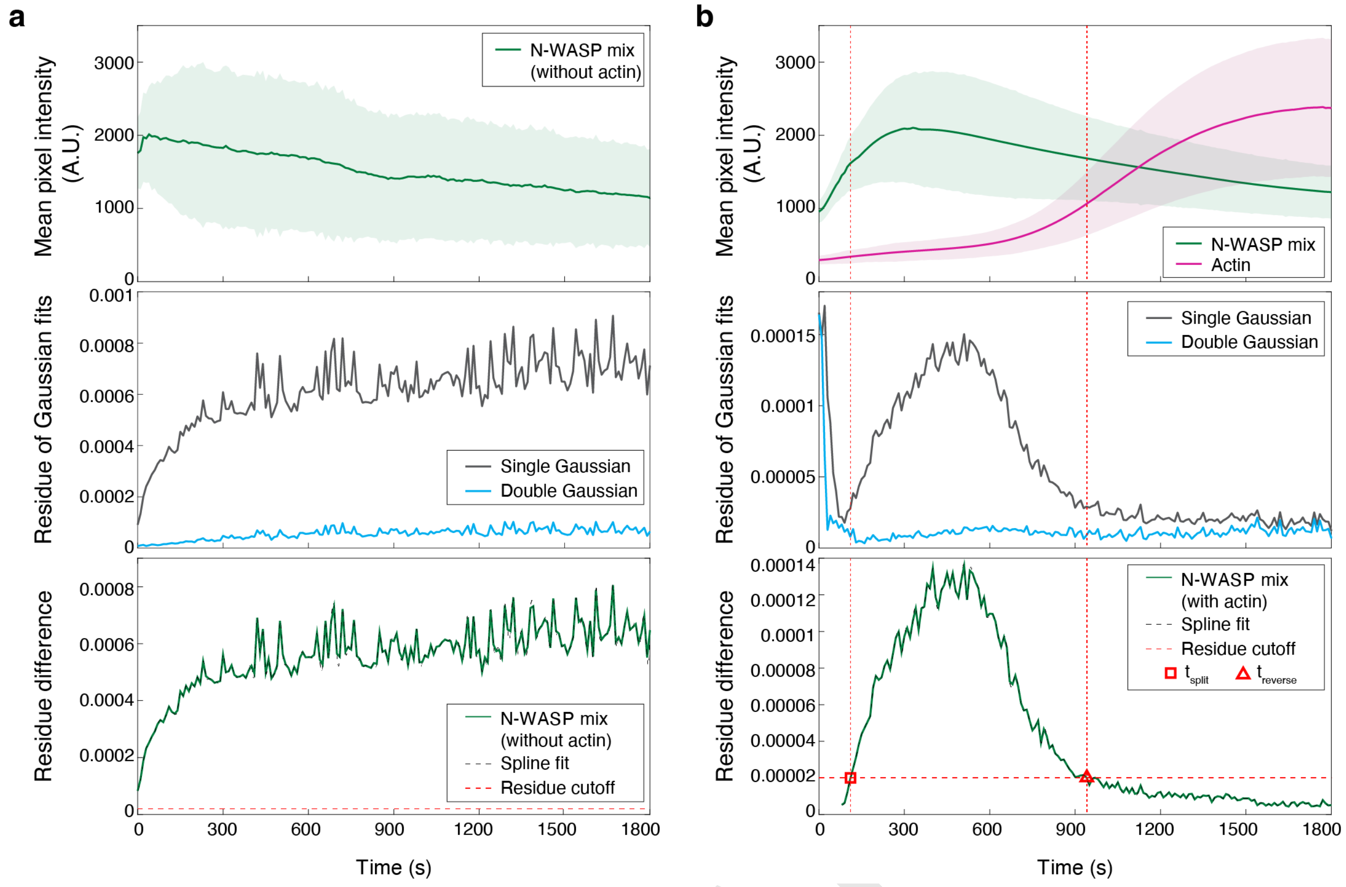


**Fig. S13. Segmentation-independent quantification of N-WASP condensation on SLBs in the accompany of actin polymerisation.** a) Upper: Fluorescence intensity kinetics for the dataset of N-WASP mix condensation in the absence of actin (Fig. 3a). Middle: Residue of single-variate and double-variate Gaussian fits as a function of time. Lower: Residue difference between single-variate and double-variate Gaussian fits as a function of time. b) Upper: Fluorescence intensity kinetics for the dataset of N-WASP mix condensation in the presence of actin and Arp2/3 (Fig. 3b). Middle: Residue of single-variate and double-variate Gaussian fits as a function of time. Lower: Residue difference between single-variate and double-variate Gaussian fits as a function of time. The residue difference were used to extract the split time $t_{split}$ (red square) and reversal time $t_{reverse}$ after setting a threshold (red dashed lines). The split takes place in actin nucleation phase and reversal in actin elongation phase (red dotted lines).

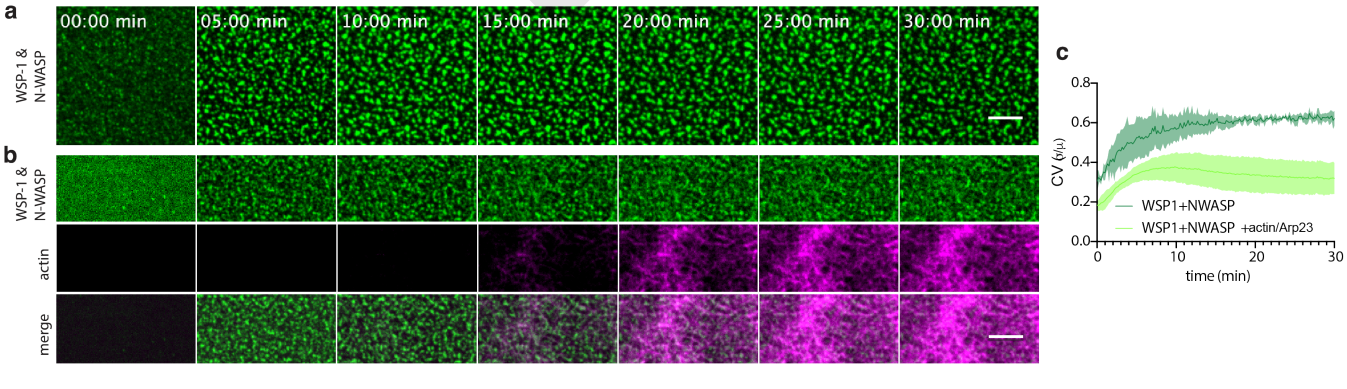


**Fig. S14. Coefficient of variation analysis of human and *C. elegans* N-WASP forming surface co-condensates in the presence of actin.** a) Confocal snapshots of a timelapse of the condensation of a binary mixture of *C. elegans* $His_6$-WSP-1 and *H. sapiens* $His_6$-N-WASP (500 nM total, 10 %AF488 labeled, green) on a SLB with 1 % Ni-NTA. b) Same as in a in the presence of actin (magenta) and Arp2/3. All scale bars = 5 $\mu$m. c) Coefficient of variation (CV) of N-WASP intensities in a field of view on the SLB over time.

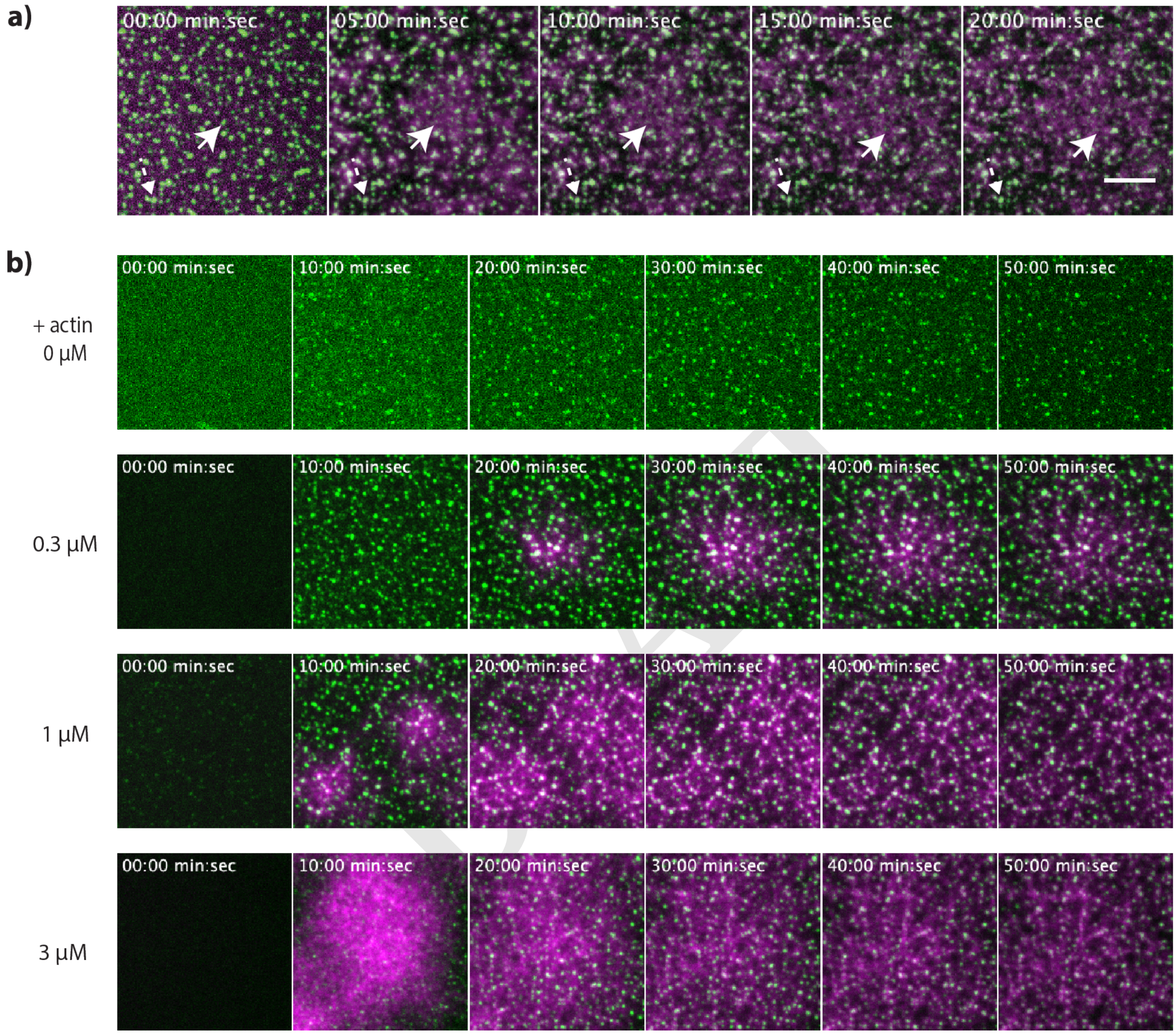


**Fig. S15. Actin polymerization disassambles N-WASP surface condensates.** a) Confocal snapshots of a timelapse (see Movie S7) of the condensation of a binary mixture of *C. elegans* $His_6$-WSP-1 and *H. sapiens* $His_6$-N-WASP (500 nM total, 10 %AF488 labeled, green) with actin (1 $\mu$M, 10 %AF647 labeled, magenta) and Arp2/3 (10 nM) on a SLB with 1 % Ni-NTA. Arrows point towards a N-WASP condensates that disassembles over time (straight line) and, where no actin polymerization is visible, condensates that grow (dotted line). b) Same as in a) with varying concentrations of actin (0-3 $\mu$M). The higher actin concentration is present in the assay, the fewer N-WASP remain in the condensed phase. Scale bars = 5 $\mu$m.

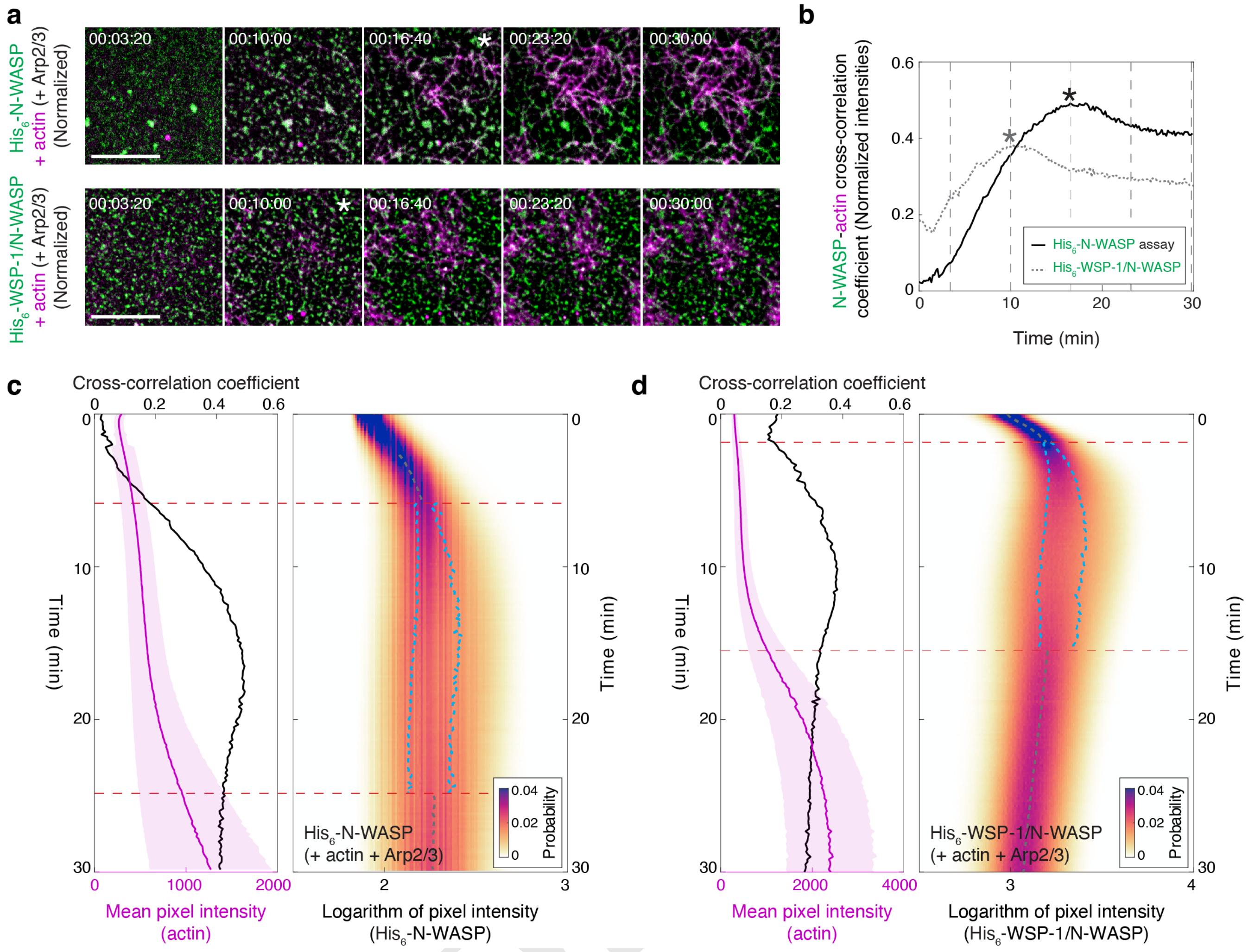


**Fig. S16. Quantification of N-WASP surface condensate disassembly in the presence of actin polymerization.** a) Confocal snapshots of two intensity-normalized timelapses of *H. sapiens* $His_6$-N-WASP (100 nM, 10 % AF488 labeled, green, same as Fig. 3f) (top row) and a binary mixture of *C. elegans* $His_6$-WSP-1 and *H. sapiens* $His_6$-N-WASP (500 nM total, 10 %AF488 labeled, green, same as Fig. 3b) (bottom row), both with actin (1 $\mu$M, 10 %AF647 labeled, magenta) and Arp2/3 (10 nM) on a SLB with 1 % Ni-NTA. Both intensities of N-WASP (green) and actin (magenta) are normalized by the mean and standard deviation of the pixel intensity distribution per snapshot (Supplementary Information). Asterisks label the snapshot with maximum pixel intensity cross-correlation coefficient (see b), also in Supplementary Information) within the five snapshots. Scale bars = 10 $\mu$m. b) Pixel intensity cross-correlation coefficient for the normalized N-WASP and actin intensity fields, calculated from the two time lapses in a) (Supplementary Information). Asterisks label the time points when the cross-correlation is maximum throughout time series. c) From the confocal timelapse of *H. sapiens* $His_6$-N-WASP with actin and Arp2/3 (Fig. 3f and a) top row), a side-by-side comparison of the dynamics of N-WASP versus actin cross-correlation coefficeint (black line) and actin mean pixel intensity (magenta line) (left panel) and the dynamics of logarithm of N-WASP pixel intensity distributions shown as kymograph (right panel). Shaded area (left panel) shows the standard deviation across the actin intensities of all pixels per snapshot. Gray line (right panel) shows the peak postion fitted from a single Gaussian while blue lines (right panel) show the peak positions fitted from a sum of two Gaussian functions. Red dashed lines (both panels) mark the split and later reversal-of-split in time. d) From the confocal timelapse of a binary mixture of *C. elegans* $His_6$-WSP-1 and *H. sapiens* $His_6$-N-WASP with actin and Arp2/3 (Fig. 3b and a) bottom row), a side-by-side comparison of the dynamics of N-WASP versus actin cross-correlation coefficeint (black line) and actin mean pixel intensity (magenta line) (left panel) and the dynamics of logarithm of N-WASP pixel intensity distributions shown as kymograph (right panel). Shaded area and colored lines in both panels represent the same quantification as in c).

**Table 1.** Protein sequences

| Protein | organism | plasmid | sequence |
|---|---|---|---|
| Untagged WSP1 | *C. elegans* | TH1828 | MSVYPPTPTMSMMNGGVDRK RAKRPPNVGSKELNSQENEM LFALVGSEAVCLTAAVVQLL KSDRGAWRVDLPHGVISLVK DYAQRAYFLRIFDILEERIV WDFKLYKAFRAQSFPQCRKL LAFEQMENGEDGVVIGLNFF SEYEAAEFKEHLERRHAQER KSTSTTRPAHPGMPIVVSSG IGSTPTRQFEITQAYGGTIR GPPMAIGGGQGITQMGTMAA APQSHTHTDGNASSSGSWFR KDKNKKKDKKSKIKKEDISN PTNFQHKAHVGWNQDSGFSN TVYDDDMDEATKNILKAAGL ESNNLNEDDKKFVKKFIAKN YDKYVSVGSLDPSQISSPLP PPIQQHPQMNQSWNQTPVRQ YKPSFPSSAPIGSGASSYST PAAPPPPTRVESHGLAPARP PPPPPSSGTRGIAPSRPLPQ APNYGTPENRPHAVPPPPPP PPPQSFGMAPISSAAPPPPP PPPPMGLPAVGAGAPPPPPP PPPSGAGGPASVLAKLPPAQ DGRSNLLAEIQAGKQLRSVQ QTADSPKSAGGDARGDVMAQ IRQGAQLKHVDAAAEQERRK STTSGAAGMGGLAGALAKAL EERRMNMGIDDTSDDDDDED DKNEWSD |
| His6-MBP-mGFP-WSP1 | *C. elegans* | TH1614 | MGSSHHHHHHSSGRMKIEEG KLVIWINGDKGYNGLAEVGK KFEKDTGIKVTVEHPDKLEE KFPQVAATGDGPDIIFWAHD RFGGYAQSGLLAEITPDKAF QDKLYPFTWDAVRYNGKLIA YPIAVEALSLIYNKDLLPNP PKTWEEIPALDKELKAKGKS ALMFNLQEPYFTWPLIAADG GYAFKYENGKYDIKDVGVDN AGAKAGLTFLVDLIKNKHMN ADTDYSIAEAAFNKGETAMT INGPWAWSNIDTSKVNYGVT VLPTFKGQPSKPFVGVLSAG INAASPNKELAKEFLENYLL TDEGLEAVNKDKPLGAVALK SYEEELVKDPRIAATMENAQ KGEIMPNIPQMSAFWYAVRT AVINAASGRQTVDEALKDAQ TNSSSNNNNNNNNNNNSSGRL EVLFQGPMVSKGEELFTGVV PILVELDGDVNGHKFSVSGE GEGDATYGKLTLKFICTTGK LPVPWPTLVTTLTYGVQCFS RYPDHMKQHDFFKSAMPEGY VQERTIFFKDDGNYKTRAEV KFEGDTLVNRIELKGIDFKE DGNILGHKLEYNYNSHNVYI MADKQKNGIKVNFKIRHNIE DGSVQLADHYQQNTPIGDGP VLLPDNHYLSTQSKLSKDPN EKRDHMVLLEFVTAAGITLG MDELYKGSSSGRENLYFQGA AA...(WSP1 sequence see above) |
| MBP-His6-WSP1 | *C. elegans* | TH2035 | MGMKIEEGKLVIWINGDKGY NGLAEVGKKFEKDTGIKVTV EHPDKLEEKFPQVAATGDGP DIIFWAHDRFGGYAQSGLLA EITPDKAFQDKLYPFTWDAV RYNGKLIAYPIAVEALSLIY NKDLLPNPPKTWEEIPALDK ELKAKGKSALMFNLQEPYFT WPLIAADGGYAFKYENGKYD IKDVGVDNAGAKAGLTFLVD LIKNKHMNADTDYSIAEAAF NKGETAMTINGPWAWSNIDT SKVNYGVTVLPTFKGQPSKP FVGVLSAGINAASPNKELAK EFLENYLLTDEGLEAVNKDK PLGAVALKSYEEELVKDPRI AATMENAQKGEIMPNIPQMS AFWYAVRTAVINAASGRQTV DEALKDAQTNSSSNNNNNNNN NNNSSGRLEVLFQGPAAAHH HHHHSSGRENLYFQGGGASG...(WSP1 sequence see above) |
| MBP-His6-N-WASP | *H. sapiens* | TH2098 | MKIEEGKLVIWINGDKGYNG LAEVGKKFEKDTGIKVTVEH PDKLEEKFPQVAATGDGPDI IFWAHDRFGGYAQSGLLAEI TPDKAFQDKLYPFTWDAVRY NGKLIAYPIAVEALSLIYNK DLLPNPPKTWEEIPALDKEL KAKGKSALMFNLQEPYFTWP LIAADGGYAFKYENGKYDIK DVGVDNAGAKAGLTFLVDLI KNKHMNADTDYSIAEAAFNK GETAMTINGPWAWSNIDTSK VNYGVTVLPTFKGQPSKPFV GVLSAGINAASPNKELAKEF LENYLLTDEGLEAVNKDKPL GAVALKSYEEELVKDPRIAA TMENAQKGEIMPNIPQMSAF WYAVRTAVINAASGRQTVDE ALKDAQTNSSSNNNNNNNNNN NSSGRLEVLFQGPAAAHHHH HHSSGRENLYFQGGGASGMS SVQQQPPPPRRVTNVGSLLL TPQENESLFTFLGKKCVTMS SAVVQLYAADRNCMWSKKCS GVACLVKDNPQRSYFLRIFD IKDGKLLWEQELYNNFVYNS PRGYFHTFAGDTCQVALNFA NEEEAKKFRKAVTDLLGRRQ RKSEKRRDPPNGPNLPMATV DIKNPEITTNRFYGPQVNNI SHTKEKKKGKAKKKRLTKAD IGTPSNFQHIGHVGWDPNTG FDLNNLDPELKNLFDMCGIS EAQLKDRETSKVIYDFIEKT GGVEAVKNELRRQAPPPPPP SRGGPPPPPPPPHNSGPPPP PARGRGAPPPPPSRAPTAAP PPPPPSRPSVAVPPPPPNRM YPPPPPALPSSAPSGPPPPP PSVLGVGPVAPPPPPPPPPP PGPPPPPGLPSDGDHQVPTT AGNKAALLDQIREGAQLKKV EQNSRPVSCSGRDALLDQIR QGIQLKSVADGQESTPPTPA PTSGIVGALMEVMQKRSKAI HSSDEDEDEDDEEDFEDDDE WED |

DRAFT

### Supplementary Note 3: Supplementary movies

**Movie S1. WSP-1 condensates coarsen and fuse in bulk**
Confocal time-lapse imaging of *C. elegans* WSP-1 (5 $\mu$M, 10% 488-tagged, MBP-tag cleaved right before experiment and KCl concentration lowered to 150 mM), in a plane close to the cover glass over the course of 150 min. Field of view 20x20 $\mu$m.

**Movie S2. WSP-1 condensates coarsen and fuse on supported lipid bilayers**
Confocal time-lapse imaging of *C. elegans* WSP-1 (100 nM, 10% 488-tagged, MBP-tag cleaved right before experiment) in actin polymerization buffer (containing 150 mM KCl), in a plane close to the supported lipid bilayer (containing 1 % Ni-NTA) over the course of 20 min. Field of view 20x20 $\mu$m.

**Movie S3. Actin polymerizes from bulk WSP1 condensates**
Confocal time-lapse imaging of *C. elegans* WSP-1 (5 $\mu$M, 10% 488-tagged, green) mixed together with actin (3 $\mu$M, 10% AF647 labeled, magenta) and Arp2/3 (100 nM) in actin polymerizing buffer containing 150 mM KCl in a plane close to the cover glass over the course of 13.5 min. Scale bar 5 $\mu$m.

**Movie S4. Actin polymerizes from WSP1 condensates on supported lipid bilayers**
Confocal time-lapse imaging of *C. elegans* WSP-1 (100 nM, 10% 488-tagged, green) mixed together with actin (1 $\mu$M, 10% AF647 labeled, magenta) and Arp2/3 (100 nM) in actin polymerizing buffer containing 150 mM KCl in a plane close to the supported lipid bilayer over the course of 20 min. Scale bar 5 $\mu$m.

**Movie S5. N-WASP adsorption and condensation on supported lipid bilayers**
Confocal time-lapse imaging of human N-WASP (10% 488-tagged, MBP-tag cleaved right before experiment, upper row 100 nM, middle 250 nM, low 500 nM) in actin polymerization buffer (containing 150 mM KCl), in a plane close to the supported lipid bilayer (containing 1 %Ni-NTA) over the course of 10 min. Scale bar 5 $\mu$m.

**Movie S6. N-WASP adsorption and condensation on supported lipid bilayers**
Confocal time-lapse imaging of binary mixture of human N-WASP and *C. elegans* WSP-1 (500 nM, 10% 488-tagged, MBP-tag cleaved right before experiment) in actin polymerization buffer (containing 150 mM KCl), in a plane close to the supported lipid bilayer (containing 1 %Ni-NTA) over the course of 30 min. Scale bar 5 $\mu$m.

**Movie S7. Actin polymerizes from N-WASP/WSP-1 condensates on supported lipid bilayers**
Confocal time-lapse imaging of binary mixture of human N-WASP and *C. elegans* WSP-1 (500 nM, 10% 488-tagged, green) mixed together with actin (1 $\mu$M, 10% AF647 labeled, magenta) and Arp2/3 (10 nM) in actin polymerizing buffer containing 150 mM KCl in a plane close to the supported lipid bilayer over the course of 30 min. Scale bar 5 $\mu$m.

DRAFT
