## Supplementary figures and images for "Actin polymerization counteracts prewetting of N-WASP on supported lipid bilayers"

### SI_Movie1.gif

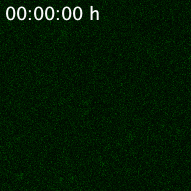

### SI_Movie2.gif

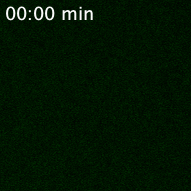

### SI_Movie3.gif

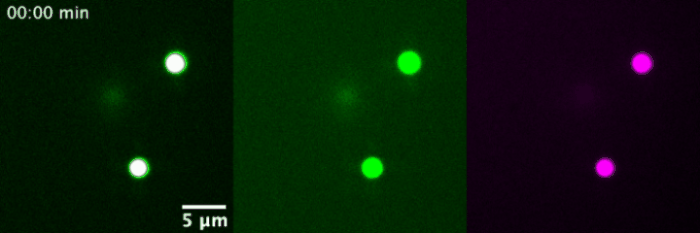

### SI_Movie4.gif

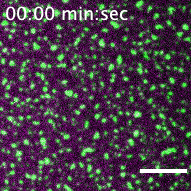

### SI_Movie5.gif

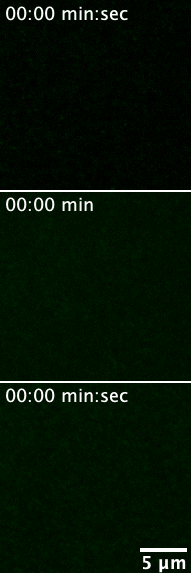

### SI_Movie6.gif

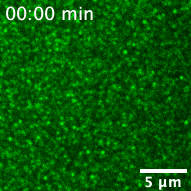

### SI_Movie7.gif

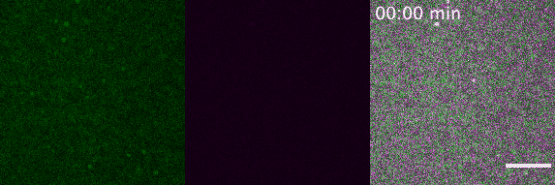
